# Nicotinic modulation of frontal circuits breaks the cycle of attentional performance failure

**DOI:** 10.64898/2026.10.05.756468

**Authors:** Tadaaki Nishioka, Kevin Norman, Yury Garkun, Justin Riceberg, Leah Waltrip, Lucy Bicks, Alesia Singh, April Serratelli, Tatsiana Mankouskaya, Priscilla Maccario, Samuel Allen, Julia Bateh, Christina Cho, Keaven Caro, Sarah Lopez, Ayako Kawatake-Kuno, Huanhuan Li, Yulong Li, Xiaoting Wu, Thomas O’loughlin, Schahram Akbarian, Hirofumi Morishita

## Abstract

The ability to recover from consecutive failures is critical for adaptive behavior, yet the underlying neural mechanisms remain unclear. *In vivo* screening of neuromodulatory inputs onto frontal-sensory cognitive control circuit identified a frontal cholinergic mechanism that counteracts lapses in attentional engagement following repeated errors. Cholinergic input from the basal forebrain to the frontal cortex was necessary and sufficient to restore performance after errors, specifically after repeated—but not after a single—failure. This effect was mediated by α7-nicotinic acetylcholine receptors in frontal-sensory cortical neurons, which sustained cortical circuit activity and frontal-sensory cortical oscillations during attentional engagement. Targeted optogenetic activation of these frontal-sensory neurons disrupted cycles of attentional performance failure and reinstated behavioral performance in a mouse model of neuropsychiatric syndrome. These findings reveal a circuit-level mechanism of cholinergic control that confers cognitive resilience under conditions of persistent failure.

## Introduction

Cognitive control enables flexible behavior by monitoring performance and initiating strategic adjustments in attentional engagement and action, particularly following errors (*1*). While isolated errors can elicit compensatory adjustments that enhance subsequent performance (*1–3*), repeated errors often lead to cumulative attentional performance failure, marked by disengagement, increased omissions, and decreased behavioral accuracy (*4–6*). The neural mechanisms that confer cognitive resilience against such maladaptive error cascades remain poorly understood.

The anterior cingulate area (ACA), a medial prefrontal cortex subregion, has been implicated in performance monitoring and attentional engagement regulation across species (*7–11*). Notably, in mice, a frontal-sensory projection from ACA to the visual cortex (VIS) —ACA_VIS_ pathway— mediates top-down attentional engagement modulation (*12–14*), especially after errors (*13, 14*), positioning this circuit as a key site for investigating the neural substrate for cognitive resilience. Here, we aimed to identify the circuit and signaling mechanisms that stabilize attentional engagement and performance in the face of persistent behavioral challenges.

We explored how error information is integrated into the frontal-sensory cognitive control circuit by focusing on upstream neuromodulatory inputs, which are known to encode feedback signals— such as errors, uncertainty, and salience—and to regulate attention engagement and performance adjustment (*1, 15–18*). To achieve this, we combined a quantitative cognitive behavior assay in mice with circuit-specific monitoring and manipulation of neural activity or neuromodulatory signaling (optogenetics, fiber photometry, and electrophysiological recordings), as well as circuit-selective gene expression profiling (RNA sequencing) and CRISPR-based gene targeting. We then leveraged these findings to restore cognitive resilience in a mouse model of neuropsychiatric syndrome.

### Cholinergic input is associated with post-error recruitment of frontal-sensory projection neurons

We first investigated the extent to which frontal-sensory ACA_VIS_ neurons exhibit trial-by-trial modulation that reflects recent error history and predicts behavioral outcomes. To achieve this, we performed projection-specific calcium imaging during the 5-choice serial reaction time task (5-CSRTT), a widely used assay for assessing attentional performance (*19–21*) (**Fig. S1**). We use the term attentional performance to refer to attentional engagement during the 5-CSRTT, operationalized primarily by the ability to accurately detect and respond to the visual stimulus while minimizing omissions. Because performance in the 5-CSRTT can also be influenced by other behavioral processes, we evaluated complementary measures, including reward collection latency, premature responding, and perseverative responding, which provide information relevant to motivation, response inhibition/impulsivity, and repetitive or compulsive-like responding, respectively (**Fig. S1**) (*19*). Thus, although the 5-CSRTT does not provide a direct, process-pure measure of attention, the primary performance measures together with complementary behavioral indices allow us to assess attentional engagement while considering other process that may contribute to task performance.

To selectively express calcium sensors in ACA_VIS_ neurons, we injected AAVrg-Cre into VIS and a Cre-dependent AAV expressing jGCaMP8s into ACA (**Fig. 1A-C, S2A**). Fiber photometry recordings revealed that ACA_VIS_ activity was modulated by recent trial outcomes, with significantly higher activity in trials where animals performed correctly after an error (E-C), compared to either consecutive correct (C-C) trials or error repeat (E-E) trials. This modulation was observed during the intertrial interval (ITI)—which reflects the anticipatory attentional engagement period— **(Fig. 1D, E)** as well as during stimulus presentation period (**Fig. S2B**). ACA_VIS_ activity was further maintained after error feedback (incorrect or omission) whereas it rapidly declined following reward delivery after correct responses (**Fig. S2C–F**). To quantify how well neural signals predict upcoming trial outcomes, we computed time-resolved area under the receiver operating characteristic curve (auROC) values comparing correct and error trials. On trials following errors— but not on trials following correct responses—auROC values gradually increased toward the end of the ITI, this indicates that ACA_VIS_ activity increasingly differentiated successful (E-C) from failed (E-E) outcomes (**Fig. 1F**) when the ACA_VIS_ activity is causally essential for post-error adjustment of attentional engagement (*14*). These results suggest that ACA_VIS_ neurons carry information relevant for adaptive control specifically after errors, when adjustments of attentional engagement are needed, and highlight this circuit as a key site for investigating upstream neural substrates.

**Fig. 1.**
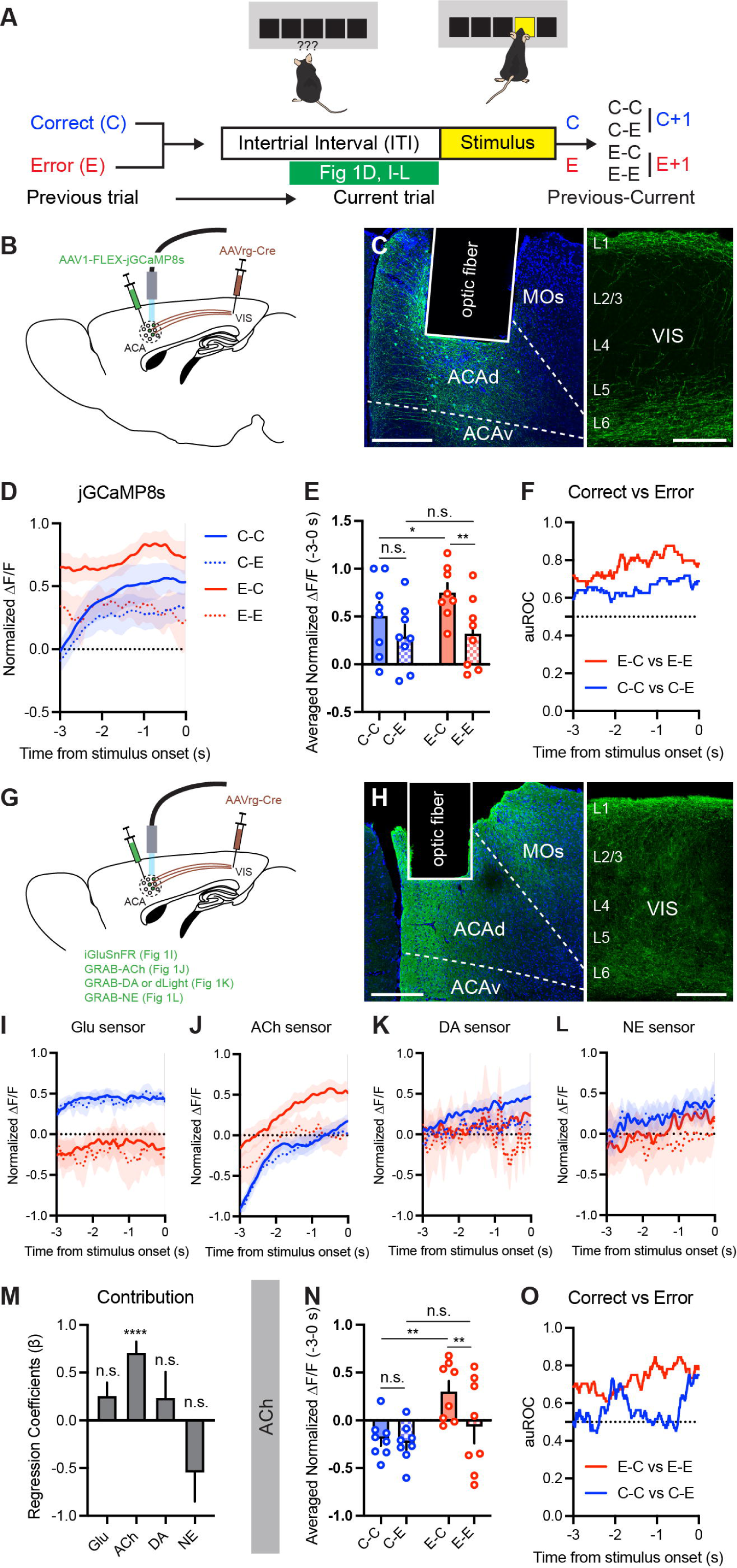
Neural dynamics of ACA_VIS_ neurons and their afferent inputs during an anticipatory period. (**A**) Schematic of trial progression in 5-CSRTT. A mouse initiates a trial with a nose poke, which begins the 5-s inter-trial interval (ITI) during which the mouse anticipates the upcoming stimulus. Trials can be analyzed in the context of past trial history on current performance, such as following a correct (C+1) or error (E+1) trial. (**B**) Surgical schematic for GCaMP imaging in the ACA_VIS_ circuit. (**C**) Expression of GCaMP in ACA (left) and VIS (right), counterstained with DAPI (blue). Scale bar, 200 µm. (**D**) jGCaMP8s signal dynamics in ACA_VIS_ neurons during the anticipatory attentional engagement period. Blue solid line: C-C trials; blue dashed: C-E; red solid: E-C; red dashed: E-E. (**E**) Averaged ACA_VIS_ activity during the anticipatory period (−3 to 0 s before stimulus onset) (Two-way RM ANOVA, effect of current trial outcome: *F*_1,7_ = 17.67, *P* = 0.0040, n = 8 mice). Post hoc comparisons (Sidak’s test) indicated that E-C activity was significantly higher than C-C (*P* = 0.038) and E-E (*P* = 0.0022), whereas C-C vs C-E (*P* = 0.062) or C-E vs E-E (*P* = 0.92) were comparable. (**F**) auROC time course for jGCaMP8s signals comparing correct and error trial types. (**G**) Schematic of viral injection sites for each biosensor. (**H**) Expression of GRAB-ACh in ACA (left) and VIS (right), counterstained with DAPI (blue). Scale bar, 200 µm. (**I–L**) Normalized fluorescence dynamics of each biosensor expressed in ACA_VIS_ neurons during the anticipatory period: (I) Glutamate (iGluSnFR, n = 7 mice), (J) Acetylcholine (GRAB-ACh, n= 8 mice), (K) Dopamine (GRAB-DA or dLight, n = 6 mice), and (L) Norepinephrine (GRAB-NE, n = 4 mice). Trial types are indicated as in panel D. (**M**) Contribution of each input to ACA_VIS_ activity during the task, assessed by regression coefficients (β). ACh input showed a significant contribution (\*\*\*\**P* < 0.0001), others were not significant (Glu, *P* = 0.087; DA, *P* = 0.40; NE, *P* = 0.090). (**N**) Mean ACh input signal during the anticipatory period (−3 to 0 s before stimulus onset) (Two-way RM ANOVA, effect of current trial outcome: *F*_1, 7_ = 10.95, *P* = 0.013). Post hoc comparisons (Sidak’s test) indicated that E-C activity was significantly higher than C-C (*P* = 0.0017) and E-E (*P* = 0.0084), whereas C-C vs C-E (*P* = 0.87) or C-E vs E-E (*P* = 0.19) were comparable. (**O**) Time course of auROC for ACh signals, comparing correct and error trials. n.s., not significant; \**P* < 0.05; \*\**P* < 0.01; \*\*\*\**P* < 0.0001. Data are presented as mean ± SEM.

To determine what upstream signals drive ACA_VIS_ activation during the late anticipatory period— when it drives post-error adjustment of attentional engagement (*14*)—we next monitored multiple neuromodulatory and glutamatergic signals targeting ACA_VIS_ neurons during the period of anticipatory attentional engagement. We selectively expressed genetically encoded sensors for glutamate (Glu), acetylcholine (ACh), dopamine (DA), and norepinephrine (NE) in ACA_VIS_ neurons(*22–26*), and performed fiber photometry recordings in mice engaged in the 5-CSRTT (**Fig. 1G, H, S3A–C, S4A**). Among these inputs, ACh activity was the only one that showed preferential elevation on trials following error responses (**Fig. 1I–L**). Specifically, while Glu signals were higher on trials following correct responses (**Fig. 1I, S3D**), ACh signals were selectively elevated on trials following errors (**Fig. 1J, N**). DA and NE signals did not differ significantly between trial types (**Fig. 1K, L, S3E, F**). The dissociation between ACA_VIS_ activity and extracellular Glu likely reflects differences between population activity and bulk extracellular Glu measurements, which integrate signals from multiple sources and clearance mechanisms.

To quantify which input best accounted for trial-by-trial variation in ACA_VIS_ activity, we performed linear regression analysis. Only ACh signals significantly predicted ACA_VIS_ activity (**Fig. 1M**), indicating a selective functional link between cholinergic input and ACA_VIS_ neuron engagement. Importantly, ACh activity was the highest on E-C trials, significantly greater than on C-C or E-E trials during both the anticipatory period (**Fig. 1N**) and stimulus presentation (**Fig. S4B**), mirroring the trial-type-dependent ACA_VIS_ activation pattern (**Fig. 1E**). ACh signals in ACA_VIS_ neurons persisted after feedback, regardless of trial type (**Fig. S4C–F**). Notably, correct response latency— a behavioral metric related to visual processing speed and attentional engagement—tended to be negatively correlated with ACh activity on E+1 trials, but not on C+1 trials (**Fig. S4G–J**). Finally, auROC analysis confirmed that ACh input most effectively distinguished correct from error outcomes on trials following error responses (**Fig. 1O**), supporting its role in encoding trial outcome in a history-dependent manner. Together, these results reinforce the view that cholinergic input is selectively associated with a control of attentional engagement following errors, in a manner consistent with the error-history-dependent activation of ACA_VIS_ neurons.

### Cholinergic input to ACA is necessary and sufficient for recovering attentional performance after repeated errors

Our correlative findings led us to test whether cholinergic input to the ACA is necessary and sufficient for maintaining attentional engagement following errors. The major cholinergic sources of cortical acetylcholine are the basal forebrain (BF) nuclei, including the horizontal limb of the diagonal band (HDB) and nucleus basalis (NB), although contributions from other sources cannot be completely excluded (*27*). Cholinergic projections from brainstem nuclei, such as the pedunculopontine tegmental nucleus (PPN) and laterodorsal tegmental nucleus (LDT), to the ACA are comparatively sparse (*28*). In addition, although ChAT-positive interneurons have been identified within the cortex, they are relatively rare (*29*). To assess necessity of cholinergic input from the BF, we injected a Cre-dependent AAV encoding halorhodopsin (eNpHR3.0) into the BF of Chat-Cre mice and implanting optic fibers above the ACA (**Fig. 2A, B, S5A**). Following training to criterion on the 5-CSRTT, mice underwent optogenetic testing. Light was delivered during the late ITI (0–3.0 s before stimulus onset) (**Fig. 2C**), a period previously shown to be critical for post-error control of attentional engagement (*14*). Initial analysis revealed a non-significant trend toward impaired performance on E+1 trials (**Fig. 2D**), prompting a more granular analysis of trial history. We therefore subdivided E+1 trials based on the outcomes of the two preceding trials: CE+1 trials, where an error trial followed a correct trial, and EE+1 trials, where two consecutive errors preceded the current trial (**Fig. 2C**). Strikingly, optogenetic inhibition impaired performance specifically on EE+1 trials (**Fig. 2E**), indicating that cholinergic input is required for maintaining performance after repeated errors. In contrast, performance on C+1 trials—whether following correct (CC+1) or error (EC+1) trials—was unaffected by ACh suppression (**Fig. 2E**), further supporting that the modulatory effect is specific to trials following repeated errors. While stimulus duration significantly influenced task performance after errors (*P* < 0.0001 (E+1), 0.0009 (EE+1)), confirming that shorter stimulus durations increased task difficulty, we found no significant interaction between stimulus duration and light condition on post-error performance (*P* = 0.5647 (E+1), 0.7519 (EE+1)), indicating that the effect of optogenetic suppression was not dependent on task difficulty. Importantly, stimulus duration was randomly varied across trials and could not be predicted by the animals. Therefore, animals were required to maintain a consistently high level of attention throughout the task, which likely minimized systematic interactions between stimulus duration and trial-history-dependent performance adjustments.

**Fig. 2.**
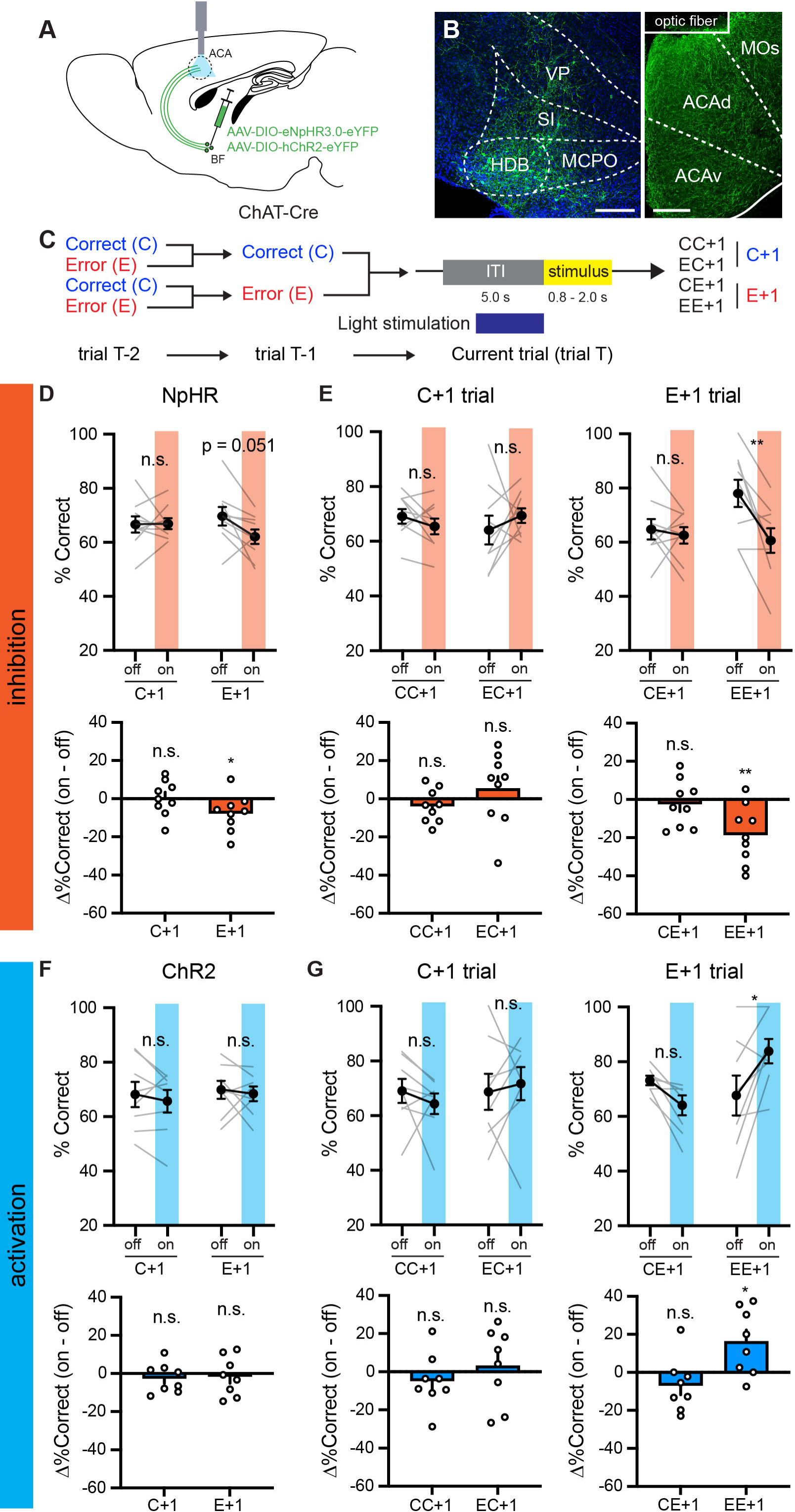
Bidirectional modulation of post-error attentional performance by BF cholinergic inputs to the ACA. (**A**) Schematic of viral injection and optic fiber implantation targeting cholinergic BF-to-ACA projections. (**B**) Expression of eNpHR3.0 in the BF (left) and ACA (right), counterstained with DAPI (blue). Scale bar, 200 µm. (**C**) Diagram of the light stimulation protocol during the 5-CSRTT in the context of consecutive trial history. Light was delivered during the ITI (-3 to 0 s before stimulus onset). (**D**) Inhibition of BF-to-ACA cholinergic projections did not significantly impair performance when analyzed based on single-trial history (Two-way RM ANOVA, effect of history: *F*_1,8_ = 0.1756, *P* = 0.69, n = 9 mice), although there was a trend toward reduced performance following error trials (E+1; *P* = 0.051, Sidak’s test. Bottom: One-sample t test, *P* = 0.047), but not following correct trials (C+1; *P* = 0.99, Sidak’s test. Bottom: One-sample t test, *P* = 0.93). (**E**) Optogenetic inhibition of BF-to-ACA cholinergic projections selectively impaired performance following consecutive errors (EE+1 trials). (Top: Two-way RM ANOVA, effect of history: *F*_1,16_ = 6.057, *P* = 0.026; off vs. on in EE+1 trials: *P* = 0.0022, Sidak’s test. Bottom: One-sample t test, *P* = 0.0069, n = 9 mice). No significant effects were observed in other trial types (CC+1, *P* = 0.72; EC+1, *P* = 0.52; CE+1, *P* = 0.87, Sidak’s test. One-sample t test, CC+1: *P* = 0.72, EC+1: *P* = 0.52, CE+1, *P* = 0.87). (**F**) Tetanic (60 Hz) stimulation of BF-to-ACA cholinergic inputs did not significantly affect performance when analyzed based on single-trial history (Two-way RM ANOVA, effect of history: *F*_1,7_ = 0.5353, *P* = 0.49; C+1, *P* = 0.65; E+1, *P* = 0.85, Sidak’s test. Bottom: One-sample t test, C+1: *P* = 0.40, E+1: *P* = 0.70). (**G**) Optogenetic activation of BF-to-ACA cholinergic projections selectively improved performance following consecutive errors (EE+1 trials). (Top: Two-way RM ANOVA, effect of history: *F*_1, 14_ = 8.376, *P* = 0.012; off vs. on in EE+1 trials: *P* = 0.024, Sidak’s test. Bottom: One-sample t test, *P* = 0.033). No significant effects were observed in other trial types (CC+1, *P* = 0.71; EC+1, *P* = 0.86; CE+1, *P* = 0.43, Sidak’s test. One-sample t test, CC+1: *P* = 0.39, EC+1: *P* = 0.68, CE+1, *P* = 0.22). n.s., not significant; \**P* < 0.05; \*\**P* < 0.01. Data are presented as mean ± SEM.

While the 5-CSRTT does not provide a direct measure of attention, the primary performance measures together with complementary behavioral indices allow us to assess attentional engagement while considering other process that may contribute to task performance (**Fig. S5B– E**). Importantly, ACh inhibition did not alter other behavioral metrics. The optogenetic inhibition did not alter reward collection latency, arguing against a major contribution from altered motivation (**Fig. S5B**). It also did not increase premature responding, arguing against altered impulsivity or response inhibition as the primary explanation **(Fig. S5D)**. Perseverative responding was also unchanged, providing no evidence that the observed phenotype was driven by repetitive or compulsive-like responding (**Fig. S5E**). Finally, all animals underwent extensive training before experimental testing, reducing the likelihood that the observed effects were attributable to failure to acquire the procedural components of the task. Collectively, these behavioral measures suggest that the observed deficit reflects an impairment in attentional engagement rather than changes in motivation or impulsivity. Trial-type analysis further indicated that the performance deficit on EE+1 trials was primarily driven by an increase in omissions (**Fig. S5F**). These results demonstrate that cholinergic input from the BF to ACA is specifically required to sustain attentional engagement following repeated errors.

To determine whether this input is also sufficient to enhance attentional performance under similar conditions, we expressed channelrhodopsin-2 (ChR2) in the BF cholinergic neurons by injecting a Cre-dependent AAV into Chat-Cre mice and implanted an optic fiber above the ACA (**Fig. 2A, S6A**). *Ex vivo* tetanic photostimulation of BF-to-ACA projections elicited inward currents in ACA_VIS_ neurons (**Fig. S6B**), confirming effective cholinergic transmission, consistent with a prior study (*30*). *In vivo* optogenetic activation during the anticipatory attentional engagement period selectively improved performance on EE+1 trials (**Fig. 2F, G**), without affecting motivation-related measures or other task variables (**Fig. S6C–F**). Trial-type analysis indicated that the performance improvement on EE+1 trials was primarily driven by decreased omissions (**Fig. S6G**). The effect of optogenetic activation was not dependent on task difficulty as we did not observe significant interaction between stimulus duration and light condition on post-error performance (*P* = 0.1715 (E+1), 0.7677 (EE+1)). A potential limitation is that optogenetic stimulation was assigned on a trial-by-trial basis with a 50% probability, and the number of EE+1 trials was relatively limited, which may introduce sampling variability in condition-level estimates. Thus, the apparent differences in EE+1 off performance between conditions may reflect stochastic trial allocation and variability in the animals contributing to each condition rather than systematic differences in the underlying off condition. However, our main conclusions are based on within-animal comparisons and remain robust to these considerations. Together, these findings demonstrate that cholinergic input from the BF to ACA is both necessary and sufficient for maintaining attentional performance specifically after repeated errors, primarily by preventing lapses in attentional engagement reflected in omission errors.

### α7-nAChRs in frontal-sensory projection neurons are required for performance recovery after repeated errors

Having established that cholinergic input from the BF to ACA is critical for recovering attentional performance after repeated errors, and given that ACA_VIS_ neurons receive direct cholinergic inputs from BF neurons (*31, 32*), we next investigated which specific acetylcholine receptor (AChR) subtypes in ACA_VIS_ neurons may mediate this effect. To identify candidate receptors, we injected AAVrg-Cre into the VIS of stop-TRAP-GFP mice, microdissected the ACA, and performed fluorescence-activated nuclear sorting (FANS) to isolate GFP+/NeuN+ double-positive nuclei, corresponding to ACA_VIS_ neurons (**Fig. 3A**). RNA sequencing of these sorted cells revealed high expression of several AChR subtypes, including *Chrm1*, *Chrm2*, *Chrm3*, *Chrna7*, and *Chrnb2* (**Fig. 3B**), as well as other neuromodulatory receptors (**Fig. S7A**). Notably, the expression profiles of these AChRs in ACA_VIS_ neurons differed from those of comparable layer 5 IT-type neurons in the anterior lateral motor cortex (ALM) and visual cortex (VIS) in the Allen Institute cell-type database (*33*) (**Fig. S7B**). Based on the observed positive correlation between ACh input and ACA_VIS_ activity (**Fig. 1**), we prioritized *Chrm1*, *Chrm3*, *Chrna7*, and *Chrnb2*—receptors with known excitatory roles—for functional testing. We designed sgRNAs targeting each receptor and assessed the efficacy of the CRISPR manipulations at the targeted genomic loci. This analysis revealed substantial indel formation and high KO scores for *Chrm1, Chrnb2, Chrna7,* with more modest disruption o *Chrm3* (**Fig.3C and Fig.S8**). We then assessed the functional consequences of receptor deletion in ACA_VIS_ neurons using a dual-virus approach: AAVrg-Cre was injected into the VIS, and a Cre-dependent AAV expressing SaCas9 and sgRNA was injected into the ACA (**Fig. 3C**, **S9A**). *Ex vivo* calcium imaging confirmed the functional effect of the knockout: carbachol-evoked responses were significantly attenuated in hemispheres with receptor deletion in ACA_VIS_ neurons (**Fig. S9B**). Following a four-week recovery, mice that had previously reached behavioral criterion on the 5-CSRTT were retrained and tested. Of the four receptor types targeted, only deletion of *Chrna7* in ACA_VIS_ neurons led to significantly lower performance on trials following errors than on trials following correct responses (**Fig. 3D**). No such effect was observed in mice with deletions of *Chrnb2*, *Chrm1*, *Chrm3*, or in control animals (**Fig. 3D**). We note that the absence of a statistically significant effect does not by itself establish that a receptor is not involved, particularly when genomic disruption is incomplete, as observed for *Chrm3*.

**Fig. 3.**
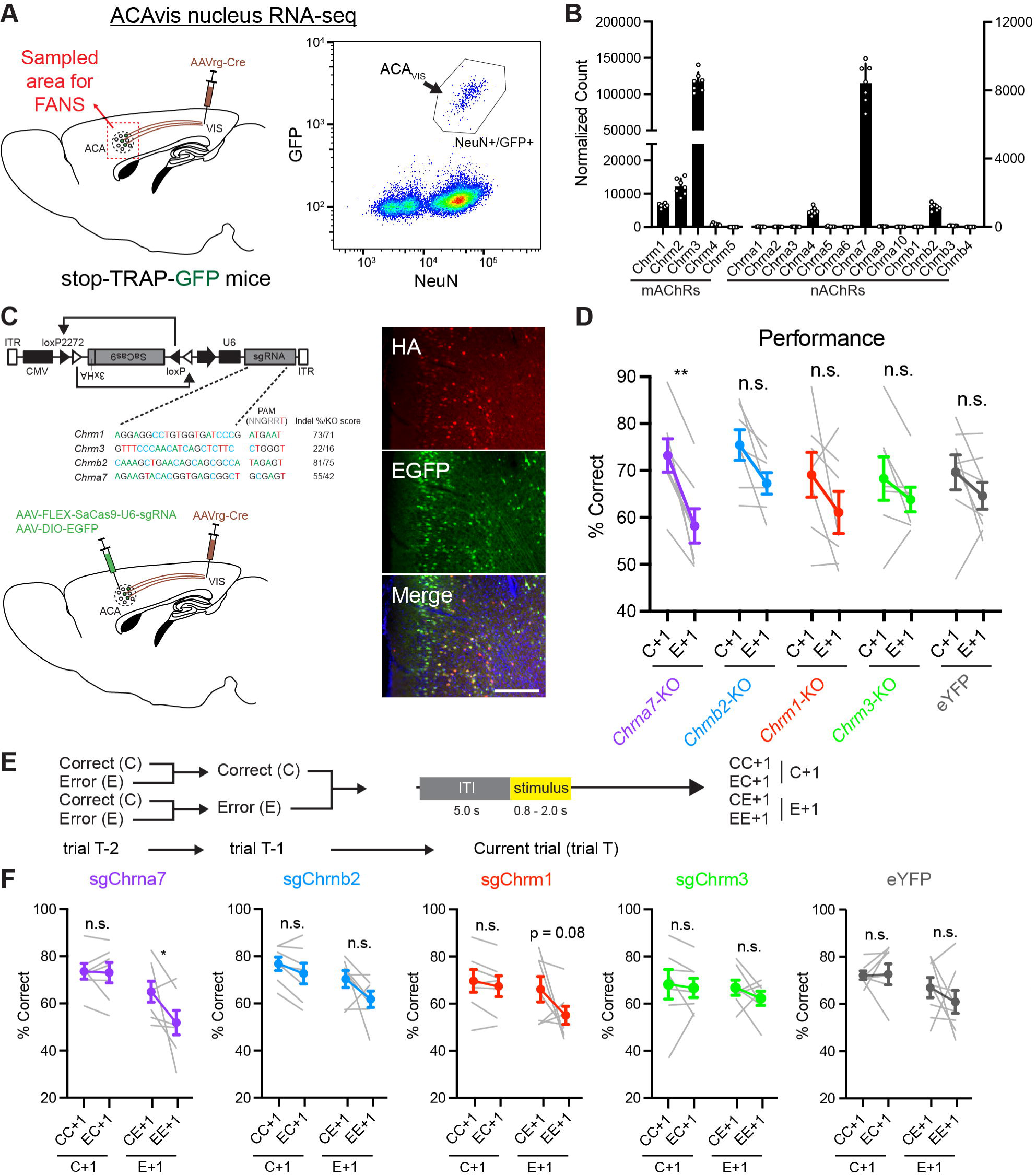
Knockdown of α7-nAChRs in ACA_VIS_ neurons impairs attentional performance following consecutive errors. **(A)** Schematic of virus injection (left) and representative FANS analysis (right) to isolate ACA_VIS_ neurons. **(B)** Expression profile of muscarinic (mAChRs) and nicotinic (nAChRs) acetylcholine receptor subtypes in ACA_VIS_ neurons, based on normalized RNA-seq read counts using DESeq2 size-factor normalization. **(C)** Design of CRISPR constructs and injection strategy (left). Efficacy of the CRISPR manipulations were assessed with ICE analysis, based on indel % and KO score. Right panels show expression of HA-tagged SaCas9 and EGFP in the ACA_VIS_ neurons following virus injection, counterstained with DAPI (blue). **(D)** Knockdown of α7-nAChR (n = 7 mice) significantly reduced performance after an error trial (E+1; Two-way ANOVA, effect of history: *F*_1,7_ = 26.07, *P* = 0.0014; *Chrna7*-KO, *P* = 0.0015, Sidak’s test), while knockdown of other AChR subtypes (β2-nAChR, n = 7 mice, M1-mAChR, n = 7 mice, M3-mAChR, n = 7 mice) or eYFP control (n = 8 mice) had no significant effects (*Chrnb2*-KO, *P* = 0.15; *Chrm*1-KO, P = 0.16; *Chrm3*-KO, *P* = 0.72; eYFP, *P* = 0.56; Sidak’s test). **(E)** Schematic diagram illustrating trial classification based on the outcome history of the previous two trials. **(F)** α7-nAChR knockdown selectively impaired performance on trials following two consecutive errors (EE+1) (Two-way ANOVA for sgChrna7, effect of history: *F*_1,12_ = 8.437, *P* = 0.013; CC+1 vs. EC+1, *P* = 0.99; CE+1 vs. EE+1, *P* = 0.038, Sidak’s test). Other knockdown groups (sgChrnb2, sgChrm1, sgChrm3) and eYFP controls showed no significant deficits (*Chrnb2*-KO, CC+1 vs. EC+1, *P* = 0.55 and CE+1 vs. EE+1, *P* = 0.11; *Chrm1*-KO, CC+1 vs. EC+1, *P* = 0.088 and CE+1 vs. EE+1, *P* = 0.085; *Chrm3*-KO, CC+1 vs. EC+1, *P* = 0.92 and CE+1 vs. EE+1, *P* = 0.50; eYFP, CC+1 vs. EC+1, *P* = 0.99 and CE+1 vs. EE+1, *P* = 0.51). n.s., not significant; \**P* < 0.05; \*\**P* < 0.01. Data are presented as mean ± SEM.

To examine whether this impairment was modulated by extended error history, we further stratified trials based on the outcomes of the two preceding trials (**Fig. 3E**). Mice with *Chrna7* deletion showed a pronounced performance deficits specifically on EE+1 trials (**Fig. 3F**). This effect was absent in all other receptor knockout groups (**Fig. 3F**), suggesting that α7-nAChRs in ACA_VIS_ neurons are preferentially required for recovering attentional performance after repeated errors. The effect of *Chrna7* deletion was not dependent on task difficulty as we did not observe significant interaction between stimulus duration and genotype condition on post-error performance (*P* = 0.5514 (E+1), 0.9468 (EE+1)). The performance deficit in the *Chrna7* knockout group was not accompanied by changes in other behavioral metrics, including reward collection latency, premature responses, or perseverative errors (**Fig. S9C–F**), indicating that the observed deficit reflects an impairment in attentional engagement. Notably, omissions increased significantly on EE+1 trials in this group (**Fig. S9G**), in line with results from optogenetic ACh inhibition and stimulation (**Fig. S5F, S6G**). Furthermore, correct response latency was significantly prolonged on EE+1 trials in *Chrna7*-knockout mice (**Fig. S9H**), echoing our observation that ACh signals in ACA_VIS_ neurons showed a trend toward a negative correlation with correct response latency on E+1 trials (**Fig. S4H, J**). Together, these results identify α7-nAChRs in ACA_VIS_ neurons as a critical molecular substrate for performance recovery after repeated errors.

### α7-nAChRs in frontal-sensory projection neurons mediate circuit activity and frontal-sensory oscillations supporting performance recovery after repeated errors

While α7-nAChRs are necessary for recovering attentional performance, it remained unclear whether they directly contribute to the enhancement of ACA_VIS_ activity observed after errors. Having established that both ACA_VIS_ activity and cholinergic input correlate with attentional performance after errors, and that cholinergic signaling via α7-nAChRs is causally involved in recovery after repeated errors, we next investigated whether α7-nAChRs in ACA_VIS_ neurons mediate the observed enhancement of ACA_VIS_ activity on the trials following errors. To address this, we conducted fiber photometry-based calcium imaging of ACA_VIS_ neurons during the 5-CSRTT in mice with or without α7-nAChR knockdown in ACA_VIS_ neurons (**Fig. 4A**, **S10A**). We first analyzed trial-type-specific activity during the anticipatory attentional engagement period (−3– 0 s before stimulus onset). While activity on C+1 trials did not differ between groups, ACA_VIS_ activity on E+1 trials was significantly reduced in α7-nAChR knockouts compared to controls, indicating a selective disruption of error-driven enhancement (**Fig. 4B, C**). To further dissect the effect of trial outcome combinations, we stratified E+1 trials by current and preceding trial types. α7-nAChR knockouts showed significantly reduced ACA_VIS_ activity on E-C trials, where a correct response followed an error, but not on E-E trials (**Fig. S10B, C**), suggesting that the α7-nAChR is specifically required to boost ACA_VIS_ activity during successful post-error recovery.

**Fig. 4.**
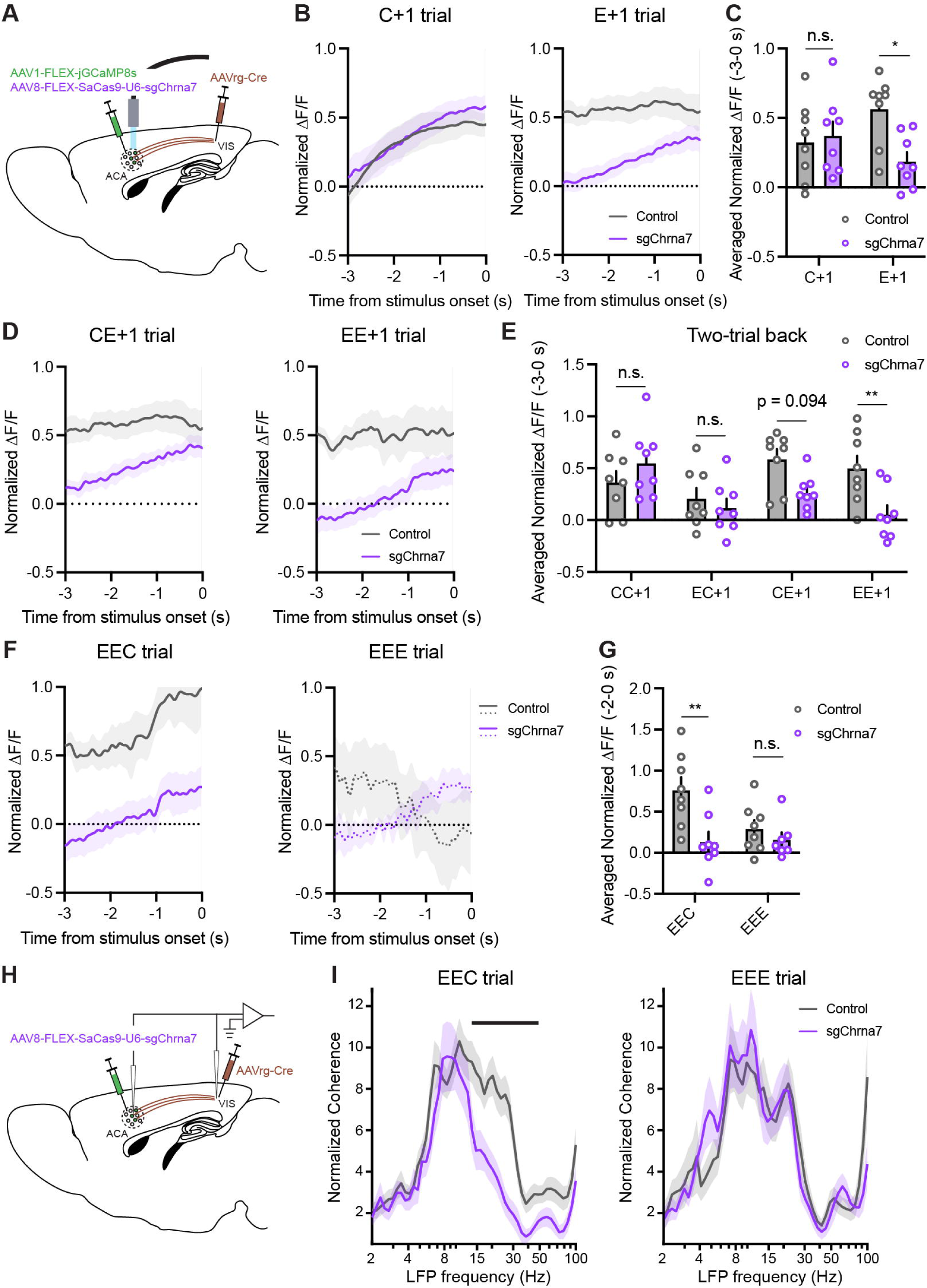
Activation of ACA_VIS_ neurons via α7-nAChRs is essential for attentional performance following consecutive errors. (**A**) Schematic of the viral injection strategy for *in vivo* calcium imaging. AAVrg-Cre and AAV1-FLEX-jGCaMP8s were injected into the VIS, and AAV8-FLEX-SaCas9-U6-sgChrna7 was injected into the ACA to knock down α7-nAChRs in ACA_VIS_ neurons. (B) jGCaMP8s signal dynamics in ACA_VIS_ neurons during the 5-CSRTT in mice with α7-nAChR knockdown (sgChrna7) (n = 8 mice). Control traces are reused from Fig. 1 (n = 8 mice). Left: Group-averaged traces for trials following correct (C+1); Right: Group-averaged traces for trials following error (E+1). Gray: control; purple: sgChrna7. (**C**) ACA_VIS_ activity during the anticipatory attentional engagement period (−3 to 0 s before stimulus onset) is significantly reduced in the sgChrna7 group compared to Control in E+1 trials (Two-way RM ANOVA, genotype × trial type interaction: *F*_3, 42_ = 5.568, *P* = 0.023; post hoc Sidak’s test: C+1, *P* = 0.92; E+1, *P* = 0.012). (**D**) jGCaMP8s signal dynamics in ACA_VIS_ neurons during E+1 trials further segmented by two-trial history. Left: Trials preceded by a correct and an error (CE+1); Right: Trials preceded by two consecutive errors (EE+1). (**E**) ACA_VIS_ activity during the anticipatory period (−3 to 0 s before stimulus onset) is significantly attenuated in sgChrna7 mice specifically in EE+1 trials (Two-way RM ANOVA, genotype × trial type interaction: *F*_3, 42_ = 5.568, *P* = 0.0026; post hoc Sidak’s test: CC+1, *P* = 0.57; CE+1, *P* = 0.95; CE+1, *P* = 0.094; EE+1, *P* = 0.0087). (**F**) jGCaMP8s signal dynamics in ACA_VIS_ neurons during the anticipatory period on EEC (left) or EEE trials (right). (**G**) ACA_VIS_ activity during the anticipatory period (−2 to 0 s before stimulus onset) in sgChrna7 mice is significantly lower compared with control mice on EEC trials but not on EEE trials (Two-way RM ANOVA, genotype × trial type interaction: *F*_1, 13_ = 6.155, *P* = 0.028; post hoc Sidak’s test: EEC, *P* = 0.0018; EEE, *P* = 0.69). (**H**) Schematic of the viral injection strategy and the LFP implantation surgery for LFP recordings in ACA and VIS during the 5-CSRTT. (**I**) Normalized ACA-VIS coherence across frequency band during anticipatory period (−3 to 0 s before stimulus onset) on EEC and EEE trials in the sgChrna7 and control group. The black line indicates frequency clusters in which the metric differs across genotypes (*P* < 0.05, cluster-based permutation test). Low-gamma coherence is significantly lower in sgChrna7 mice compared to control mice on EEC trials (sgChrna7: 43 trials from 3 mice, control: 64 trials from 3 mice) but not on EEE trials (sgChrna7: 27 trials from 3 mice, control: 35 trials from 3 mice). n.s., not significant; \**P* < 0.05; \*\**P* < 0.01. Data are presented as mean ± SEM.

We next examined whether trial history extending two trials back modulated this activity. Although reductions in ACA_VIS_ activity were observed on both CE+1 and EE+1 trials in knockouts, only the reduction on EE+1 trials reached significance (**Fig. 4D, E**). The effect of α7-nAChRs knockout on suppressing ACA_VIS_ activity was specific to the anticipatory period preceding correct responses, but not error responses following repeated errors (**Fig. 4F, G**). We then assessed whether the number of consecutive prior errors affected the ability of ACA_VIS_ activity to predict behavioral performance. In control mice, correct responses were associated with significantly higher ACA_VIS_ activity than errors, but only after two consecutive errors—not after one or three (**Fig. S11A, B**). This effect was abolished in α7-nAChR knockouts (**Fig. S11C, D**). Together, these findings demonstrate that α7-nAChRs are essential for error-driven enhancement of frontal-sensory ACA_VIS_ neuron activity, particularly after two consecutive errors—when transient compensatory mechanisms begin to fail.

We next investigated the extent to which α7-nAChRs in ACA_VIS_ neurons influence network-level oscillatory dynamics between ACA and VIS in performance recovery after repeated errors. To this end, we performed local field potential (LFP) recordings from both ACA and VIS during the 5-CSRTT in mice with or without α7-nAChR knockout in ACA_VIS_ neurons (**Fig. 4H**). Although α7-nAChR knockout did not affect LFP power in ACA or VIS during correct or error trials following repeated errors (EEC or EEE) (**Fig. S12A-C**), it significantly reduced ACA-VIS coherence in the low-gamma frequency range during the anticipatory period leading to correct responses but not error responses after two consecutive errors (**Fig. 4I**). Phase-locking value (PLV), a measure of phase synchrony between ACA and VIS, was similarly decreased in the low-gamma band on EEC trials but not on EEE trials (**Fig. S12D**). Collectively, these findings indicate that α7-nAChRs in ACA_VIS_ neurons are critical for promoting long-range coordination of ACA-VIS oscillations following repeated errors that support performance recovery.

### Optogenetic stimulation of frontal–sensory projection neurons rescues the attentional performance decline after repeated errors in a mouse model of neuropsychiatric syndrome

While activation of ACA_VIS_ neurons via α7-nAChRs is crucial for maintaining attentional performance following repeated errors in wild-type (WT) mice, how abnormalities in this mechanism contribute to the cognitive impairments associated with neuropsychiatric disorders remains unclear. The chromosomal 15q13.3 microdeletion, a 1.5 megabase region encompassing 6 genes including *CHRNA7* gene encoding α7-nAChR, is one of three major copy number variants strongly associated with a subset of individuals diagnosed with psychiatric disorders such as autism and schizophrenia(*34, 35*). Notably, mice heterozygous for this deletion (15q13.3/+) exhibit attentional performance deficits (*36, 37*). To examine whether these mice display a history-dependent decline in attentional engagement, we analyzed their performance on the 5-CSRTT. The 15q13.3/+ mice displayed a selective decrease in performance on trials following repeated errors (**Fig. 5A**), with performance significantly lower than that of WT controls on those trials (**Fig. 5B**). Other behavioral parameters, including response latencies and impulsivity measures, were comparable between genotypes (**Fig. S13A–D**), indicating that the observed deficit reflects an impairment in attentional engagement following repeated errors.

**Fig. 5.**
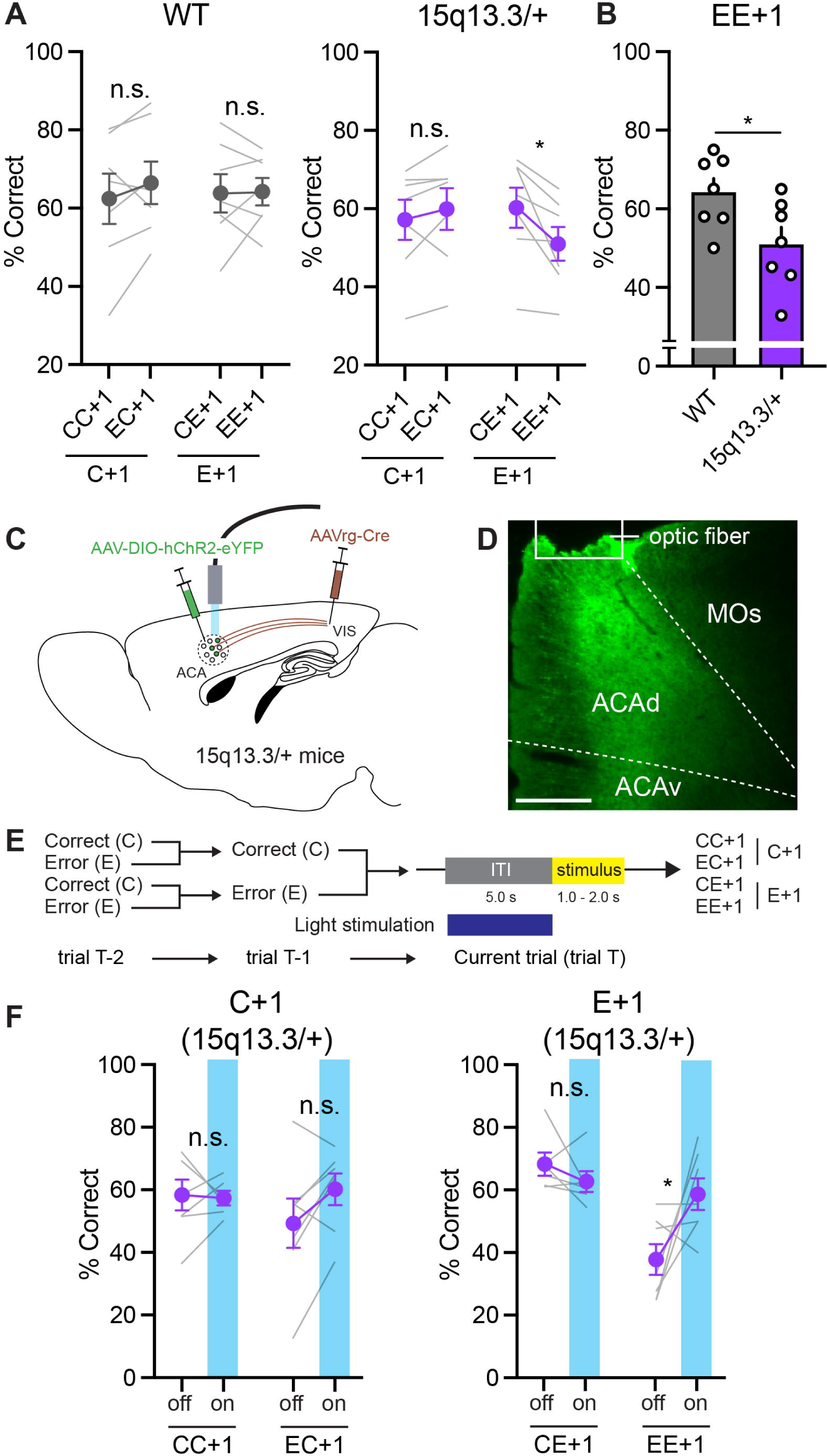
Optogenetic activation of ACA_VIS_ neurons normalizes attentional performance deficits following repeated errors in 15q13.3/+ mice. **(A)** While WT mice showed no difference across trial types, 15q13.3/+ mice exhibited a selective performance deficit specifically in trials following repeated errors (Two-way RM ANOVA for WT, trial type × trial type interaction: *F*_1,12_ = 0.6632, *P* = 0.43; CC+1 vs. EC+1, *P* = 0.39; CE+1 vs. EE+1, *P* = 0.99, Sidak’s test; Two-way RM ANOVA for 15q13.3/+, trial type × trial type interaction: *F*_1,12_ = 9.128, *P* = 0.011; CC+1 vs. EC+1, *P* = 0.58; CE+1 vs. EE+1, *P* = 0.013, Sidak’s test, n = 7 mice per group). (**B**) Performance on EE+1 trials was significantly lower in 15q13.3/+ mice compared with WT mice (Unpaired t-test, *P* = 0.035). **(C)** Schematic showing viral injection and optic fiber implantation targeting ACA_VIS_ neurons in 15q13.3/+ mice. **(D)** Representative image of ChR2 expression in ACA_VIS_ neurons. Scale bar, 200 µm. (**E**) Schematic of the optogenetic stimulation protocol during the 5-CSRTT, aligned to consecutive trial history. Light stimulation (473 nm, 30 Hz) was delivered during the inter-trial interval (ITI). (**F**) Optogenetic activation of ACA_VIS_ neurons selectively improved performance on EE+1 trials in 15q13.3/+ mice (Two-way RM ANOVA, light × trial type interaction: *F*₁,_4_ = 11.81, *P* = 0.026; off vs. on in EE+1 trials: *P* = 0.013, Sidak’s test, n = 7 mice). No significant effects were observed in other trial types (CC+1, *P* = 0.95; EC+1, *P* = 0.089; CE+1, *P* = 0.59; Sidak’s test). n.s., not significant; \**P* < 0.05. Data are presented as mean ± SEM.

We next examined whether optogenetic activation of ACA_VIS_ neurons could rescue this impairment. Trained 15q13.3/+ mice received bilateral injections of a Cre-dependent AAV encoding ChR2 in the ACA and an AAVrg-Cre in the VIS, followed by implantation of optical fibers above the ACA (**Fig. 5C**). This strategy enabled selective expression of ChR2 in ACA_VIS_ neurons (**Fig. 5D, S13E**). After retraining, optical stimulation was delivered during the anticipatory attentional engagement period of the 5-CSRTT to activate ACA_VIS_ neurons (**Fig. 5E**). Both our prior stimulation findings (*14*) and our current ACA-VIS coherence results (**Fig. 4I**) suggest that 30 Hz (low-gamma frequency) stimulation may improve performance. We thus optogenetically stimulated ACA_VIS_ neurons at 30 Hz during the anticipatory period and found that blue light illumination selectively improved performance on trials following repeated errors (**Fig. 5F**), without affecting other trial types or behavioral parameters (**Fig. S13F–I**). Together, these results identify an attentional performance deficit following repeated errors in 15q13.3/+ mice and demonstrate that activation of the ACA_VIS_ circuit is sufficient to restore attentional performance after repeated errors.

## Discussion

In this study, we identified a frontal cholinergic mechanism that selectively supports recovery from attentional performance decline following repeated errors. Our findings support a model in which increased cholinergic input from the BF boosts frontal-sensory ACA_VIS_ activity and ACA-VIS oscillations after errors via α7-nAChRs, thereby preventing lapses in attentional performance. This mechanism is especially critical after repeated errors, when the risk of falling into a maladaptive error loop is highest. By driving α7-nAChRs-mediated neural engagement, the system may actively disrupt cycles of attentional disengagement and behavioral failure (**Fig. 6**). Whereas prior studies have focused on immediate post-error effects—such as post-error slowing, often attributed to increased caution or transient motor inhibition (*1*)—our findings uncover a distinct, history-dependent control mechanism recruited under cumulative failure. Recovery from repeated errors does not reflect a simple amplification of the single-error response, but instead engages a qualitatively different slower-acting regime of cognitive control. The selective emergence of the circuit effect after two consecutive errors suggests a nonlinear relationship between error history and adjusting attentional engagement. A single error may produce a modest perturbation that can be readily compensated for, whereas two consecutive errors may engage a cholinergic-dependent adjustment detectable in our assay. The absence of a comparable effect after three errors further suggests that this relationship is not monotonic and may involve additional limiting processes with greater error accumulation. These possibilities remain to be tested directly.

**Fig. 6.**
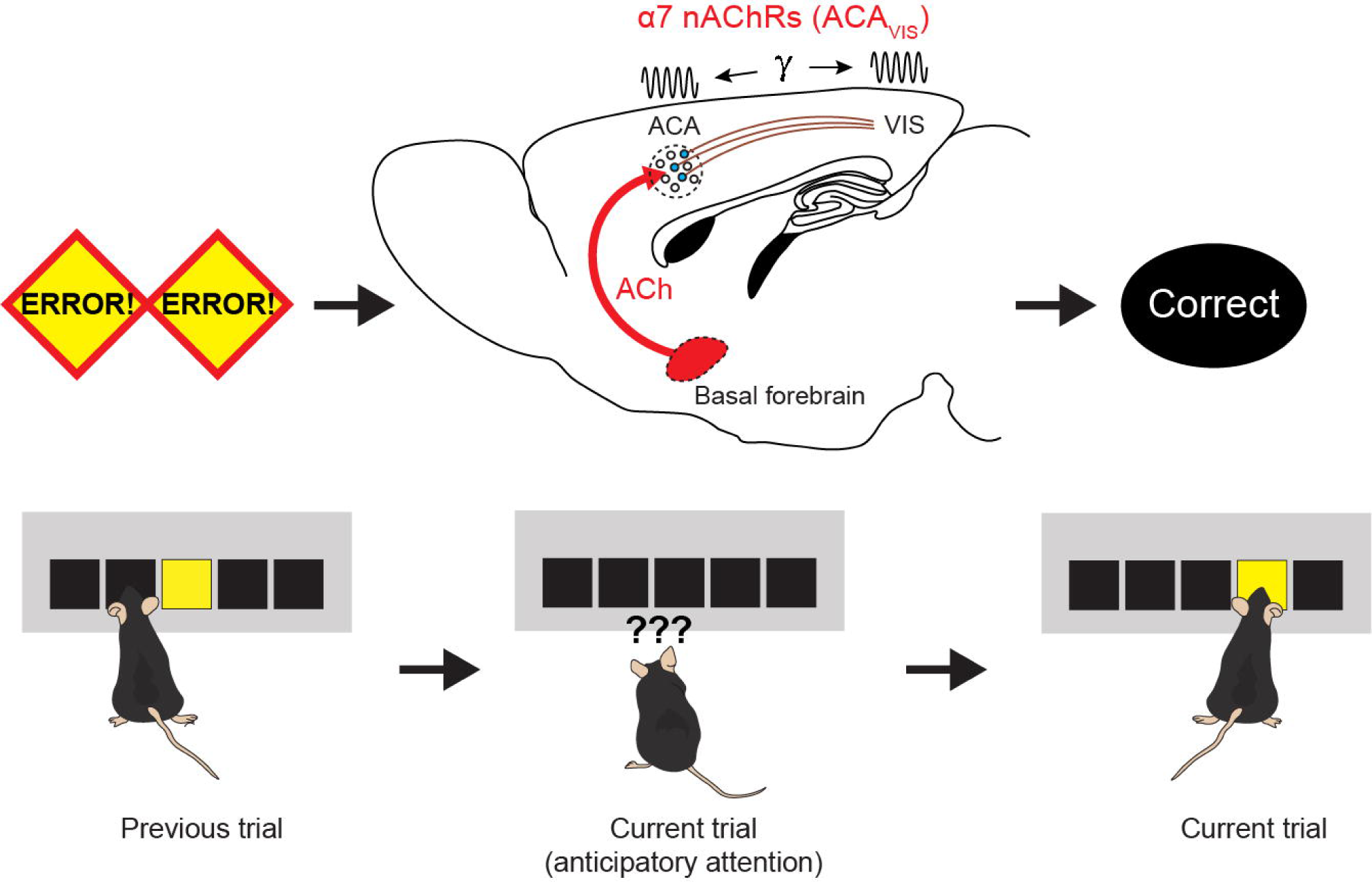
Schematic circuit model for overcoming lapses in attentional engagement after repeated errors. In this model, after consecutive errors, ACh released from the basal forebrain modulates frontal-sensory ACA_VIS_ projection neurons and frontal-sensory cortical oscillations via α7-nAChRs. This activation is critical for disrupting a negative cycle in which reduced attentional engagement following errors leads to further errors.

Such error-history dependence could allow the brain to scale cognitive-control resources according to the reliability and significance of performance feedback, avoiding excessive recruitment in response to isolated errors while enabling stronger compensation when errors indicate persistent performance deterioration. Thus, the multiple-error dependence observed here may reflect an adaptive feature rather than a limitation of the mechanism. Importantly, the threshold for engaging this mechanism is likely context dependent: a single error with greater salience, uncertainty, or aversive consequence could potentially generate a sufficiently strong control signal. Accordingly, the number of preceding errors may represent one determinant of the relevant cognitive state rather than an obligatory trigger for cholinergic recruitment. Testing whether a single highly salient error can engage the same mechanism by manipulating error significance or outcome value will be an important direction for future studies.

Our findings support a model in which the ACh system dynamically modulates performance across multiple timescales, depending on task demands and error history. Prior studies have shown that the ACh system is involved in both fast and slow performance adjustments across different contexts (*38–43*). In the domain of attention, while ACh release has not been linked to attention on very short (*41*) or slow timescales (*44*), chronic manipulations of the ACh system support its role in sustaining attentional performance (*45–47*). Our study may help reconcile these seemingly disparate findings by identifying a context at an intermediate timescale—specifically, during trials following consecutive errors—when ACh signaling adjusts attentional engagement levels to promote behavioral recovery. This interpretation is consistent with theoretical and empirical models suggesting that animals assign errors to disturbances operating at distinct timescales to guide learning and adaptation (*48, 49*). While other neuromodulatory systems such as DA or NE did not exhibit error-history-dependent modulation during the anticipatory period in our task (Fig.1G-M), they may nevertheless contribute to behavioral adaptation on different timescales or in different contexts, particularly given that frontal–sensory cortical neurons also express receptors for neuromodulators beyond ACh (**Fig. S7A**). DA may support strategic behavioral adjustments over longer timescales via basal ganglia circuits, potentially relaying history information in a delayed or indirect manner (*50, 51*). NE may contribute to more global, gradual changes in cognitive resource allocation or fatigue, shaping performance over extended periods and in distinct behavioral contexts (*52, 53*).

Post-error enhancement of frontal cholinergic signaling may prevent lapses in attentional engagement by modulating cortical network dynamics through frontal-sensory projections. Two key hallmarks of attention—reduced local noise correlation (*54*) and enhanced long-range cortical oscillation (*55*) —are both known to be promoted by ACh (*18*). Notably, the ACh-induced desynchronized cortical state (*56, 57*), characterized by reduced low-frequency local field potential activity, has been associated with post-error correction driven by ACA_VIS_ modulation (*14*). Supporting this, a recent study shows that desynchronized cortical activity is selectively important for perceptual decision making following behavioral errors (*58*). ACh is also known to enhance long-range coherence between cortical regions (*59, 60*)—another key neural correlate of attention (*55, 61, 62*)—driven in part by projections from the frontal cortex to sensory areas (*63*). This is consistent with our findings that frontal-sensory cortical ACh signaling drives low-gamma frontal-sensory oscillations following repeated errors, which support performance. Our LFP findings further suggest that α7-nAChR-dependent recruitment of ACA_VIS_ neurons may enhance coordination between ACA and VIS following repeated errors, potentially facilitating communication between frontal cognitive-control signals and sensory representations. Although the correlational nature of the LFP measurements precludes concluding that ACA–VIS synchrony itself causally restores attention, this coordination may represent a candidate circuit mechanism for post-error attentional adjustment and re-engagement of sensory processing. Together, these findings position cholinergic modulation of ACA_VIS_ neurons as a dynamic, history-sensitive mechanism that may help prevent lapses in attentional engagement by promoting coordinated frontal-sensory network dynamics and enabling context-dependent adjustments in visual attention.

More broadly, the cholinergic modulation of ACA_VIS_ neurons may represent one example of a broader principle in which cholinergic signaling modulates top-down frontal interactions with sensory cortical areas, including frontal–auditory or frontal–somatosensory pathways, where cholinergic modulation could similarly regulate the re-engagement of task-relevant sensory processing following attentional disruption (*64*). Such mechanisms would not necessarily be identical across sensory modalities, as their contribution may depend on the behavioral context, sensory modality, task demands, and nature of the error. We therefore do not interpret the present findings as demonstrating that the ACA_VIS_ pathway is uniquely responsible for breaking the cycle of attentional failure. Determining whether similar cholinergic mechanisms operate across other frontal–sensory pathways will require direct circuit-level investigation.

Among the various AChR subtypes, our study highlighted α7-nAChRs as key modulators of ACA_VIS_ neurons and recovery of attentional engagement after repeated errors. Although the role of α7-nAChRs in attention has been debated (*65, 66*), our findings suggest that their contribution within frontal circuits may become particularly important under conditions of high attentional demand, such as after repeated behavioral failures. This framework is further supported by findings from a mouse model of a neuropsychiatric syndrome (15q13.3/+) characterized by reduced expression of α7-nAChRs, which displays attentional performance deficits selectively after repeated errors—deficits that can be rescued by stimulating ACA_VIS_ neurons. Although other cholinergic receptor subtypes, including β2-nAChRs (*65*) and M1-AChRs (*67–69*), may contribute in distinct contexts, our data do not establish their involvement; however, nonsignificant effects do not exclude a contribution, particularly with incomplete receptor disruption. Consistent with this possibility, previous human pharmacological work demonstrated that muscarinic signaling contributes to post-error modulation of visual cortical processing and behavioral adjustment (*68*), supporting a broader role for acetylcholine in adaptive attentional control. Thus, while our findings specifically implicate α7-nAChRs, contributions from other cholinergic receptor subtypes or functional redundancy among them remain possible. Targeting the identified frontal cholinergic pathways and α7-nAChRs may offer therapeutic entry points for enhancing cognitive resilience (*70, 71*), particularly in neuropsychiatric conditions characterized by deficits in cognitive control (*72–74*).

## Materials and Methods

### Animals

Adult male C57BL/6 mice (Charles River Laboratories, MA), Chat-Cre mice (#006410, Jackson Laboratory), Cre-dependent eGFP-L10a mice (#024750, Jackson Laboratory), and 15q13.3/+ mice (gifted by Lundebeck) were used in this study. Mice were group-housed in a temperature- and humidity-controlled vivarium under a standard 12-hour light/dark cycle, with all behavioral experiments conducted during the light phase. Training began when the mice were 8–10 weeks old. During behavioral training, mice were water-restricted and given access to water for 2 hours per day, while their body weight was maintained at approximately 85–90% of their free-feeding weight. Food was available ad libitum throughout the experiment. All experimental procedures were approved by the Institutional Animal Care and Use Committee at the Icahn School of Medicine at Mount Sinai.

### Stereotaxic surgery

Stereotaxic surgeries were conducted according to procedures described in previous studies(*14, 75*). Mice were anesthetized with 2% isoflurane and positioned in a stereotaxic apparatus (Narishige, East Meadow, NY). For circuit-specific calcium imaging of ACA_VIS_ neurons, injection coordinates were based on previous studies(*14, 75*). AAV1-Syn-FLEX-jGCaMP8s (Addgene) was injected into the anterior cingulate area (ACA), and AAVrg-EF1a-Cre (Boston Children’s Research Viral Core) was concurrently injected unilaterally into the visual cortex (VIS). For ACA injections, coordinates relative to bregma were: AP +0.2 mm, ML 0.4 mm, DV −0.6 & −1.0 mm. For VIS injections, coordinates relative to lambda were: AP 0.0 mm, ML 2.85 mm, DV −0.5 mm; AP 0.0 mm, ML 3.15 mm, DV −0.5 mm; and AP +0.3 mm, ML 3.0 mm, DV −0.5 mm. Injection sides were counterbalanced. A fiber-optic ferrule (1.3 mm length, 0.48 NA, 400 µm core diameter; Doric Lenses, Quebec, Canada) was implanted at the center of the ACA (AP +0.2 mm, ML 0.4 mm, DV −0.8 mm) and secured to the skull using dental cement (C&B Metabond, Parkell, Edgewood, NY; Jet Acrylic, Lang Dental, Wheeling, IL). For input imaging experiments, AAV5-Syn-FLEX-iGluSnFR.A184S (Addgene), AAV8-EF1a-DIO-GRAB-gACh4l (Yulong Li lab), AAV1-CAG-DIO-dLight3.6 (Neurotools), or AAV8-Syn-DIO-GRAB-DA3m (Yulong Li lab) was injected into the ACA instead of AAV1-Syn-FLEX-jGCaMP8s (Addgene). For norepinephrine input imaging, AAVrg-GRAB-NE1m (Addgene) was injected into VIS. In these cases, the fiber-optic ferrule was implanted at coordinates: AP +0.2 mm, ML 0.4 mm, DV −0.8 mm. Injection coordinates for basal forebrain (BF) targeting were based on previous studies(*41, 64*). For optogenetic manipulations of cholinergic projections, AAV2-EF1a-DIO-eNpHR3.0-eYFP (UNC vector core) or AAV2-EF1a-DIO-hChR2 (H134R)-eYFP (UNC vector core) was bidirectionally injected into the BF of heterozygous Chat-Cre mice. BF injection coordinates relative to bregma were: AP +0.2 mm, ML ±1.3 mm, DV −4.5 & −5.5 mm; AP −0.2 mm, ML ±1.3 mm, DV −4.5 & −5.5 mm; and AP −0.9 mm, ML ±2.2 mm, DV −3.3 & −5.0 mm. In both eNpHR3.0 and ChR2 experiments, bilateral 470 nm LEDs (500 µm diameter; Amuza, San Diego, CA) were implanted at the ACA (AP +0.2 mm, ML ±0.5 mm, DV −0.7 mm). For optogenetic stimulation of ACA_VIS_ neurons, AAV2-EF1a-DIO-hChR2 (H134R)-eYFP (UNC vector core) was bidirectionally injected into ACA and AAVrg-EF1a-Cre (Boston Children’s Research Viral Core) into VIS of 15q13.3/+ mice. Bilateral 470 nm LEDs (500 µm diameter; Amuza, San Diego, CA) were implanted at the ACA (AP +0.2 mm, ML ±0.5 mm, DV −0.7 mm). For the transcriptome study, AAVrg-EF1a-Cre was injected into the VIS of Cre-dependent eGFP-L10a mice at the following coordinates relative to lambda: AP 0.0 mm, ML 2.85 mm, DV −0.5 mm; AP 0.0 mm, ML 3.15 mm, DV −0.5 mm; and AP +0.3 mm, ML 3.0 mm, DV −0.5 mm. For *ex vivo* validation of CRISPR-based viral vector systems, rAAV-EF1a-Cre was injected bilaterally into the VIS, while AAV1-hSyn-FLEX-jGCaMP8s was injected into the left ACA and 1:1 mixture of AAV1-hSyn-FLEX-jGCaMP8s and the CRISPR-based viral vector was injected into the right ACA. For CRISPR-based acetylcholine receptor knock-down behavioral experiments, either AAV8-FLEX-SaCas9-U6-sgChrna7, AAV8-FLEX-SaCas9-U6-sgChrnb2, AAV8-FLEX-SaCas9-U6-sgChrm1, or AAV8-FLEX-SaCas9-U6-sgChrm3 was injected into the ACA at three sites: AP +0.2 mm, ML ±0.4 mm, DV −0.7 mm; AP +0.7 mm, ML ±0.4 mm, DV −0.7 mm; and AP −0.3 mm, ML ±0.4 mm, DV −0.7 mm. Simultaneously, AAVrg-EF1a-Cre was injected into the VIS at the following coordinates relative to lambda: AP 0.0 mm, ML 2.85 mm, DV −0.5 mm; AP 0.0 mm, ML 3.15 mm, DV −0.5 mm; and AP +0.3 mm, ML 3.0 mm, DV −0.5 mm. For CRISPR combined with calcium imaging of ACA_VIS_ neurons, AAV8-FLEX-SaCas9-U6-sgChrna7 was injected into the ACA using the same three coordinates as above. AAVrg-EF1a-Cre was injected into the VIS at the same coordinates, and AAV1-Syn-FLEX-jGCaMP8s was additionally injected into the ACA at AP +0.2 mm, ML 0.4 mm, DV −0.6 & −1.0 mm. A fiber-optic ferrule (1.3 mm length, 0.48 NA, 400 µm core diameter) was implanted at AP +0.2 mm, ML 0.4 mm, DV −0.8 mm and secured using dental cement. All viral infusions (500 nL) were performed at a rate of 100 nL/min using the Nanoject III with glass-pulled syringes. The syringe was held in place for 1 minute after each injection to minimize viral backflow. Behavioral testing began at least three weeks following viral injection to allow for optimal expression.

### Behavior

#### Five-choice serial reaction time task (5-CSRTT)

##### Apparatus

Training and testing were conducted in a Bussey–Saksida touchscreen chamber (Lafayette Instruments, Lafayette, IN), housed within a sound-attenuating box. The chamber consisted of a black plastic trapezoid (walls 20 cm high; 18 cm wide at the screen-magazine side; 24 cm wide at the screen side; 6 cm wide at the magazine side). A 12.1-inch touchscreen (resolution 600 × 800) was positioned on one wall and covered by a black plastic mask with five square windows (4.0 × 4.0 cm, spaced 1.0 cm apart and positioned 1.5 cm above the floor). ABET II and WhiskerServer software (Lafayette) were used to control operant system and data collection.

##### Habituation

Mice were first habituated to the chamber and trained to retrieve diluted (30%) sweetened condensed chocolate milk (Eagle Brand, Borden, Richfield, OH) from the magazine. During this phase, the magazine light was illuminated and milk was delivered every 10 s upon tray entry. Mice were required to collect 20 rewards within two consecutive 30-min sessions to proceed. In subsequent shaping sessions, a white square stimulus appeared randomly in one of the five windows and remained on screen until touched. Correct responses triggered milk delivery, a tone, and magazine light illumination. Touches to non-stimulus windows had no programmed consequence. Mice progressed once they made at least 20 correct touches in 30 minutes across two consecutive days.

##### Basic training

Mice were tested 5–6 days per week, with each session consisting of 100 trials or lasting up to 30 minutes. Each trial began with illumination of the magazine light. Once the mouse exited the magazine, a fixed intertrial interval (ITI) was imposed before stimulus presentation. The ITI duration was adjusted according to the stimulus duration (SD), such that trials with longer SDs of 32 s and 16 s were preceded by shorter ITIs of 2 s and 3 s, respectively, whereas shorter SDs of 8 s and below were paired with a 5-s ITI. If a mouse touched the screen during the ITI, the response was recorded as premature and punished with a 5-s time-out in which the house light was turned on. A white square stimulus was then presented in one of five windows for a given SD, followed by a 5-s limited-hold period during which the stimulus disappeared but responses were still accepted. A response to the correct location was rewarded with a tone, chocolate milk delivery, and illumination of the magazine light. Responses to other windows were recorded as incorrect, and failure to respond during the stimulus or limited-hold period was considered an omission. Both incorrect responses and omissions were punished with a 5-s time-out. Repeated responses after a correct or incorrect touch, before reward collection, were recorded as perseverative responses. Training began with a 32-s SD and progressed sequentially through 16, 8, 4, and 2 s. Advancement to the next stage required at least 50 trials, at least 80% accuracy, and no more than 20% omissions for two consecutive days at each SD. After reaching these criteria at the 2-s SD for two consecutive days, mice proceeded to testing.

##### 5-CSRTT testing

To assess attentional performance under increased demand, mice were tested using reduced and pseudo-randomly shuffled SDs of 2.0, 1.5, 1.0, and 0.8 s across sessions. All experimental testing, including fiber photometry, optogenetic manipulations, and CRISPR manipulations was conducted under this condition. Between testing days, mice were retrained using a fixed 2-s SD to confirm stable baseline performance. Behavioral performance was evaluated using several parameters. Percentage correct was calculated as (100 × correct responses)/(correct responses + incorrect responses + omissions), capturing overall task engagement and attentional capacity. Percentage accuracy was defined as (100 × correct responses)/(correct responses + incorrect responses), providing a measure of choice precision independent of omissions. Percentage omission was calculated as (100 × omissions)/(omissions + correct responses + incorrect responses), reflecting inattention or disengagement. Premature response rate was defined as the number of responses during the ITI divided by the total number of trials, and served as a measure of impulsivity. Perseverative responses were defined as additional touches made to any screen following a correct response but before reward retrieval, reflecting compulsive tendencies. Correct response latency was measured as the time interval between stimulus onset and the animal’s correct touch, serving as an indicator of attentional processing speed. Reward collection latency was calculated as the time between a correct response and the animal’s entry into the reward magazine, and was used as a proxy for motivation.

##### 5-CSRTT in mice with CRISPR-mediated gene editing

Mice resumed behavioral training four weeks after surgery, beginning at SD 32 s and advancing through SDs of 16, 8, 4, and 2 s. Once animals consistently achieved ≥50 trials within a 30-minute session at SD 2 s for two consecutive days, they proceeded to CRISPR testing, which was conducted across three daily sessions. Behavioral performance during CRISPR testing was quantified as the average across these three sessions.

#### Fiber photometry recording

Mice were first implanted with a mono-fiber optic cannula (MFC_400/430-0.48_1.3mm_MF1.25_FLT, Doric Lenses) stereotaxically positioned above the ACA. Behavioral training resumed either one week (standard group) or four weeks (CRISPR group) after surgery, beginning from the 32-s stimulus duration (SD) and advancing through SDs of 16, 8, 4, and 2 s. Mice were required to achieve at least 50 trials within a 30-minute session at SD 2 s to proceed. Next, mice were habituated to the recording condition by attaching a fiber optic patch cable (Doric Lenses, MFP_400/430/0.48_1.3). Animals then repeated the sequence of SDs—32, 16, 8, 4, and 2 s—with the cable connected. Once they again achieved ≥50 trials within 30 minutes at SD 2 s for two consecutive days with the patch cable attached, they proceeded to fiber photometry testing. All photometry recordings were conducted at least three weeks after viral injection to ensure sufficient expression. On the day of recording, behavioral testing began with SD 4 s, and once mice achieved 8–12 correct responses, the session transitioned to variable SDs (2.0, 1.5, 1.0, and 0.8 s), presented in a pseudo-random sequence. The patch cable was connected to the implanted cannula using bronze sleeves, following previously described procedures(*14*). Excitation light was delivered from two LEDs (Tucker-Davis Technologies, Alachua, FL): 465 nm to excite jGCaMP8s in a calcium-dependent manner and 405 nm as an isosbestic control reference. The 465 nm and 405 nm LEDs were sinusoidally modulated at 210 Hz and 330 Hz, respectively. Light from both LEDs was coupled into a fluorescence MiniCube (Doric Lenses), where it passed through the appropriate excitation filters and was directed to the implanted optic fiber via a patch cable. Fluorescence emission returned through the same fiber and exited the MiniCube through the green emission port (500–540 nm), which transmits emission signals resulting from both excitation wavelengths. The 465 nm- and 405 nm-evoked components were separated using real-time lock-in demodulation based on their distinct modulation frequencies. Signal acquisition and demodulation were performed using a digital signal processor (RZ10X, Tucker-Davis Technologies) and the Synapse Software Suite. The final signal was sampled at 1018 Hz. The light power at the fiber tip was approximately 20 μW. The 405 nm excitation wavelength served as an isosbestic reference and was useful for evaluating potential motion-related fluctuations during recording. ΔF/F was calculated using a running median approach as follows: (470 nm signal – median 470 nm signal over the last 100 s) / (median 470 nm signal over the last 100 s), based on procedures previously described(*76*). The resulting ΔF/F signals were z-scored for all subsequent analyses. Fiber photometry data were analyzed across different trial outcomes (correct, incorrect, omission) as well as response latencies, including correct response latency (CRL), incorrect response latency (IRL), and reward collection latency (RCL). Each animal underwent two recording sessions, and data from both sessions were combined for analysis.

#### Optogenetics during 5-CSRTT

Following the same retraining protocol as used for fiber photometry, mice resumed behavioral training one week after surgery, beginning at SD 32 s and advancing through SDs of 16, 8, 4, and 2 s. Once animals consistently achieved ≥50 trials within a 30-minute session at SD 2 s, they proceeded to the headstage habituation stage. During habituation, a 470 nm LED headstage (Amuza, San Diego, CA) was connected. Animals then repeated the sequence of SDs—32, 16, 8, 4, and 2 s—while the headstage was attached. When animals again achieved ≥50 trials at SD 2 s for two consecutive days with the headstage connected, they underwent two optogenetic testing sessions on the following two days (one session per day). All optogenetic testing sessions were conducted at least three weeks after viral injection to ensure sufficient expression. On each testing day, behavioral sessions began at SD 4 s, and once mice achieved 8–12 correct responses, the session transitioned to variable SDs (2.0, 1.5, 1.0, and 0.8 s), presented in a pseudo-random sequence. Optogenetic inhibition of cholinergic input in the ACA was achieved via bilateral delivery of continuous 470 nm light using the headstage, based on previous findings that red-shifted light frequencies caused off-target effects reducing 5-CSRTT performance in control mice (White et al., 2018). Optogenetic activation of cholinergic input was performed via bilateral delivery of 60 Hz, 470 nm light using the headstage. Illumination occurred during 50% of trials, pseudo-randomly interleaved with non-light trials, and was delivered during the ITI (−3 to 0 s before stimulus onset). Each optogenetics session began with a brief warm-up phase at SD 4 s, and after achieving 8–12 correct responses, stimulus durations were pseudo-randomly varied among 2.0, 1.5, 1.0, and 0.8 s (variable SD condition), consistent with the photometry experiments. Optogenetic activation of ACA_VIS_ neurons was performed via bilateral delivery of 30 Hz, 470 nm light using the headstage. Illumination occurred during 50% of trials, pseudo-randomly interleaved with non-light trials, and was delivered during the ITI (−5 to 0 s before stimulus onset). Stimulus durations were pseudo-randomly varied between 2.0 and 1.0 s (variable SD condition). Light delivery was controlled via TTL pulses sent from the behavioral control system to a signal generator (DG1022A, Rigol Technologies), which triggered the light with a 5% duty cycle. Light output at the fiber tip was calibrated to 1–5 mW.

#### *Ex vivo* electrophysiology

Acute brain slices were prepared using a vibrating blade microtome (Leica VT1200S) in ice-cold, oxygenated artificial cerebrospinal fluid (ACSF) containing (in mM): 210.3 sucrose, 11 glucose, 2.5 KCl, 1 NaH₂PO₄, 26.2 NaHCO₃, 0.5 CaCl₂, and 4 MgCl₂. 300-μm-thick coronal slices were allowed to recover for 20 min at room temperature in cerebrospinal solution with reduced sucrose (105.2 mM) and addition of NaCl (59.5 mM). Following recovery, slices were maintained at room temperature in standard ACSF of the following composition (in mM): 119 NaCl, 2.5 KCl, 1 NaH_2_PO_4_, 26.2 NaHCO_3_, 11 glucose, 2 CaCl_2_, and 2 MgCl_2_. Somatic patch clamp recordings were performed from fluorescently labeled neurons. For patch clamp recordings, borosilicate glass electrodes (5–7 MΩ) were filled with the internal solution containing (in mM): 120 Cs-methanesulfonate, 10 HEPES, 0.5 EGTA, 8 NaCl, 4 Mg-ATP, 1 QX-314, 10 Na-phosphocreatine, and 0.4 Na-GTP (pH 7.25, 295 mOsm). Optogenetic recordings were performed in CSF (5 mM KCl, 0.5 mM MgCl₂, 1.0 mM CaCl₂), as this composition more closely resembles interstitial CSF in situ (Fishman, 1992; Sanchez-Vives & McCormick, 2000). Recordings were made at −70 mV. Signals were low-pass filtered at 1 kHz and sampled at 20 kHz. Neurons were included in the analysis if input resistance, series resistance and membrane potential did not change more than 10% during the course of recordings. For optogenetic stimulation of cholinergic synaptic inputs, ChR2-expressing terminals were stimulated with trains of blue light pulses (450 nm) generated by a high-power LED light source (Prizmatix, Israel), controlled by a Master-9 pulse generator, and delivered through a 40× water-immersion objective. Stimulation parameters were matched to those used in behavioral experiments: 0.8-ms pulses delivered at 60 Hz for 3 s. Following the optogenetic stimulation of the basal forebrain terminals in ACA, a non-specific nicotinic receptor antagonist Mecamylamine (10 μM, Tocris) was bath-applied to confirm that the recorded responses were mediated by nicotinic acetylcholine receptors. For analysis, ChR2-evoked postsynaptic currents were averaged across five repetitions and quantified as the difference between the local peak current and baseline.

#### Transcriptome analysis

##### Nuclei Isolation and Fluorescence Activated Nuclei Sorting (FANS)

After more than 3 weeks from AAV injections, Cre-dependent eGFP-L10a mice were lightly anesthetized with isoflurane and rapidly decapitated. Brains were removed and frozen. Frontal cortex tissue was dissected and then dounced in hypotonic lysis buffer, followed by ultracentrifugation through a sucrose gradient to obtain nuclei. Nuclei were resuspended in PBS with 0.01% BSA and stained with NeuN (1:1000, Millipore, MAB377A5) and DAPI (1:2000 Sigma Aldrich, 10236276001 Roche), a nuclear counter stain, prior to fluorescence activated nuclei sorting (FANS) to separate GFP+ ACAvis nuclei from GFP-non-ACAvis nuclei using an Aurora CS cell sorter (Cytek). ACAvis nuclei were collected in Trizol LS and frozen for RNA extraction (ThermoFisher, 10296028) for a final volume ratio of 3:1 Trizol LS:nuclei PBS.

##### RNA extraction, libraries, and sequencing

TRIzol/nuclei suspensions were thoroughly mixed with an equal volume of 100% ethanol and loaded onto a Zymo-Spin IC Column for RNA extraction using the Direct-zol RNA MicroPrep (Zymo, R2060) using manufacturer’s instructions. RNA was treated with DNase I for 15 minutes at room temperature. Total nuclear RNA libraries were generated using the SMARTer Stranded RNA-Seq Kit (Takara, 634836). Briefly, extracted RNA was fragmented by incubation for 4 minutes at 94°C, then stranded Oligos, RNase Inhibitor, dNTP mix, DTT, and Reverse transcriptase were incubated with the RNA at 42°C for 90 min to allow for cDNA synthesis, followed by 70°C for 10 min to terminate the reaction. cDNA was purified using SPRI AMPure XP beads (Beckman NC9959336) and then amplified with 14 cycles of PCR using the Illumina indexing primer set, followed by a final SPRI AMPure XP purification. Libraries were assessed for quality on the bioanalyzer using Agilent High Sensitivity DNA kit (Agilent 5067-4626) as well as by using Qubit. Libraries were then pooled and processed for 75 base pair paired-end sequencing by the New York Genome Center on a NovaSeq 6000 platform. Paired end Fastq files were assessed for quality using the FastQC module, and adapters were removed using Cutadapt. Trimmed fastq files were aligned to the mouse mm10 genome using the HiSAT2 aligner, duplicates were removed, and featureCounts (subread/1.5.2) was used to assign reads to genes. We quantified gene expression using both exonic and intronic reads using DESeq2 size-factor normalization to account for differences in library size across the biological replicates.

#### AAV vector construction

The full sequences of *Chrna7*, *Chrnb2*, *Chrm1*, and *Chrm3* were obtained from NCBI (https://www.ncbi.nlm.nih.gov/). To identify protospacer adjacent motif (PAM) sites and design single guide RNAs (sgRNAs), the exon sequences of these genes were analyzed using the CRISPOR web tool (https://crispor.tefor.net). Optimal sgRNA target sites were selected based on a comprehensive evaluation of their specificity scores, predicted editing efficiencies, potential off-target effects, and positions within coding exons(*77*). The following oligonucleotide pairs were designed to encode the selected sgRNA target sequences:

For *Chrna7*: 5’-CACCGAGAAGTACACGGTGAGCGGCT and 5’-AAACAGCCGCTCACCGTGTACTTCTC
For *Chrnb2*: 5’-CACCGCAAAGCTGAACAGCAGCGCCA and 5’-AAACTGGCGCTGCTGTTCAGCTTTGC
For *Chrm1*: 5’-CACCGAGGAGGCCTGTGGTGATCCCG and 5’-AAACCGGGATCACCACAGGCCTCCTC
For *Chrm3*: 5’-CACCGGTTTCCCAACATCAGCTCTTC and 5’-AAACGAAGAGCTGATGTTGGGAAACC

All oligonucleotides were synthesized with 5’ phosphorylation by Thermo Fisher Scientific and resuspended to a final concentration of 100 µM. The plasmid pAAV-FLEX-SaCas9-sgRNA (Addgene plasmid #124844) was digested with BsaI-HFv2 (New England Biolabs) for 2 hours and purified using the QIAquick Gel Extraction Kit (QIAGEN). To prepare annealed oligos, 1 µl of each complementary strand was combined with 2 µl of 5× T4 DNA Ligase buffer and 4 µl of nuclease-free water, and the mixture was subjected to gradual cooling from 100°C to 35°C in 5°C decrements every 5 minutes. After annealing, 1 µl of T4 DNA Ligase (Invitrogen) and 1 µl of the digested plasmid were added, and the ligation reaction was incubated at room temperature for 2 hours. Five microliters of the ligation mixture were transformed into E. coli Stbl3 competent cells and plated on lysogeny broth (LB) agar containing ampicillin. Following overnight incubation at 37°C, plasmids were extracted using the QIAprep Miniprep Kit (QIAGEN). Positive clones were identified by restriction digestion with SmaI. To confirm correct insertion of the sgRNA sequence, Sanger sequencing was performed using the primers 5’-GACTATCATATGCTTACCGT (forward) and 5’-CTCGCCAACAAGTTGACGAGATAAACAC (reverse). Sequencing was conducted by Genewiz. Verified plasmids were sent to Charles River Laboratories for large-scale preparation and packaging into AAV8 particles.

#### *In vitro* validation of CRISPR-mediated gene editing

##### Neuro2A cell culture and transfection

Neuro2A cells were maintained in DMEM supplemented with GlutaMAX, 10% fetal bovine serum, 50 U/mL penicillin, and 50 µg/mL streptomycin at 37°C in a humidified atmosphere containing 5% CO_2_. Cells were seeded in 6-well plates at a density of 2.5 × 10⁵ cells per well and transfected the following day using Lipofectamine 3000 (Thermo Fisher Scientific). Each well was co-transfected with 0.5 µg of AAV-EF1a-Cre-IRES-mCherry plasmid and 2.0 µg of either AAV-CAG-FLEX-EGFP or one of the following CRISPR plasmids: AAV-FLEX-SaCas9-U6-sgChrna7, AAV-FLEX-SaCas9-U6-sgChrnb2, AAV-FLEX-SaCas9-U6-sgChrm1, or AAV-FLEX-SaCas9-U6-sgChrm3. The AAV-CAG-FLEX-EGFP condition served as the unedited control. For each well, the plasmids were combined with 5 µL of P3000 reagent in 125 µL of Opti-MEM. Separately, 3.75 µL of Lipofectamine 3000 was diluted in 125 µL of Opti-MEM. The two solutions were combined and incubated for 10–15 min at room temperature before being added dropwise to the cells. Five days after transfection, cells were washed once with DPBS and collected in 800 µL of DNA/RNA Shield (Zymo Research) per well.

##### Sanger sequencing and indel analysis

Genomic DNA was extracted using the Quick-DNA Miniprep Plus Kit (Zymo Research; D4074) according to the manufacturer’s instructions. Briefly, 400 µL of each DNA/RNA Shield sample was incubated with 20 µL of proteinase K for 10 min at 55°C, mixed with an equal volume of Genomic Binding Buffer, applied to a spin column, washed, and eluted. Genomic regions encompassing the CRISPR target sites were amplified using Q5 High-Fidelity 2× Master Mix (New England Biolabs; M0492). Each 100-µL PCR reaction contained 50 µL of 2× Master Mix, 0.5 µL each of 100 µM forward and reverse primers, and 10–30 ng of genomic DNA. PCR amplification was performed with an initial denaturation at 98°C for 30 s, followed by 30 cycles of 98°C for 10 s and 72°C for 40 s, and a final extension at 72°C for 7 min. The following primer pairs were used to amplify genomic regions encompassing the respective CRISPR target sites:

For *Chrna7*: 5’-CTGCTGCACGGTAACCTCAGCG and 5’-GGAGTGAGGGCAGGAGTGGG AA
For *Chrnb2*: 5’-AGGCCCAGTCAAAGGTCCCCTC and 5’-GCATTGCCTTGTTCTGGCCCCT
For *Chrm1*: 5’-CGTGGTGCACAGACATACGCGT and 5’-TGACAGGGAGGTAGAAGGCG GC
For *Chrm3*: 5’-ACAGCCAAGTGTGGTGTGTGCC and 5’-GCAGGAGCCCACAGGACAAA GG

PCR products were verified by agarose gel electrophoresis and purified using the QIAquick PCR Purification Kit (QIAGEN; 28104). Purified amplicons were subjected to Sanger sequencing by Genewiz using the following primers:

For *Chrna7*: 5’-CGCCCACAACTGTCGTCTCCAC
For *Chrnb2*: 5’-ACCCTCTTGCCATCTCAGAAGCC
For *Chrm1*: 5’-CAGAGGTCACAGGCCAGTGTGC
For *Chrm3*: 5’-CTGCGCAGGCCAGGCTTAAGAG

Sequencing chromatograms obtained from cells transfected with the respective AAV-FLEX-SaCas9-U6-sgRNA plasmids were compared with chromatograms of the corresponding genomic loci obtained from the common unedited control sample. Indel formation was quantified by indel % and KO score using ICE CRISPR Analysis software (version 4.2; EditCo Bio). Indel % measures the total percentage of targeted genomes that have any modification (insertion or deletion) compared to the unedited control. KO score is calculated by summing the relative frequencies of all predicted indels that are expected to cause functional protein knockout (a frameshift mutation or deletions larger than 21 bp).

#### *Ex vivo* calcium imaging

For the validation of CRISPR-based viral vector systems, *ex vivo* calcium imaging was performed. Brain slices were prepared as described for in vitro electrophysiology recordings. Following recovery, slices were placed in CSF with 5 mM KCl, 0.5 mM MgCl₂, 1.0 mM CaCl₂ to mimic interstitial CSF. Slices were visualized under an epifluorescence upright microscope (Olympus BX51, Olympus, Japan) equipped with a high-power LED-coupled light source used for GCaMP8 activation (Prizmatix). The excitation light was projected onto a dichroic mirror and reflected toward the objective lens to illuminate the slice. GCaMP8 fluorescence from the slice was transmitted through an absorption filter (λ = 505 nm) to a CCD camera (DAGE-MTI, USA) connected to a PC. A water-immersion 10× objective with a 0.5× lens attached to the c-mount of the CCD camera was used to visually identify the region for imaging. The acquired images had a resolution of 2560 × 1920 pixels and a framerate of 20 Hz. To activate cholinergic receptors, carbachol (4 mM) was dissolved in ACSF and loaded into a glass pipette connected to a Picospritzer II (Parker Instrumentation, Chicago, IL). The glass pipette was positioned parallel to the midline with the tip of the pipette at the top of the slice to ensure even diffusion of carbachol to activate both hemispheres. A pulse of carbachol (150 ms) was applied using a pressure of 10 PSI. Analysis of the GCaMP fluorescence was performed in ImageJ. To allow for comparisons between control and the hemisphere with CRISPR-based viral vector expression, calcium signals were measured as the change in fluorescence relative to the baseline and normalized to the initial background measured prior to drug application (ΔF/F). Single regions of interest (ROIs) per hemisphere were selected to analyze the time course of activation.

#### LFP recordings

Mice were implanted with two 0.1-inch screws (Pinnacle; #8403) bilaterally above the cerebellum to serve as ground and common reference for LFP recording. Three single-ended Pt/Ir electrodes (Pinnacle; #8425-PR-3) were placed as follows: Electrode 1 in right VIS (AP +0.1 mm, ML +3.0 mm, DV −0.5 mm) and Electrodes 2 and 3 in bilateral ACA (Electrode 2: AP +0.2 mm, ML +0.4 mm, DV −0.8 mm; Electrode 3: AP +0.2 mm, ML −0.4 mm, DV −0.8 mm). Following placement, electrodes were secured with cyanoacrylate (Loctite) and fully covered with cement. Connections from the screws and electrodes were soldered to a 6-pin head mount (Pinnacle; #8235-SM) and encased in cement. During 5-CSRTT testing, LFP signals were captured via a tethered system and acquired using Sirenia Acquisition software. Data acquisition was performed at 2 kHz using a Pinnacle preamplifier with high-pass (0.5 Hz) and low-pass (200 Hz) filters, and a smoothing low-pass filter (6 Hz).

Time–frequency power, phase-locking, and coherence in the final 3 s of the ITI were computed using complex Morlet wavelets (6–24 cycles, scaled by frequency). For each trial, analytic signals were obtained via FFT-based convolution. Power was quantified as the squared magnitude of the wavelet coefficients. Phase-locking value (PLV) was computed as the magnitude of the circular mean of instantaneous phase differences between regions across time within each trial. Magnitude-squared coherence was estimated from within-trial cross-spectral and auto-spectral averages. Power was normalized using a session-level null distribution, created by drawing 1000 random 3-s segments from the same recording session and computing a global surrogate mean and variance per frequency. In contrast, PLV and coherence were normalized using trial-specific surrogate nulls (*78*): for each trial, 500 independent circular time-shifts were applied to one region’s signal, preserving its spectral structure while disrupting phase alignment with the other region. These trial-wise surrogate distributions provided frequency-specific means and variances for Z-scoring observed PLV and coherence. Group differences across frequencies were assessed using a one-dimensional cluster-based permutation test, which identifies significant frequency clusters by comparing observed cluster mass against a permutation-derived null distribution

#### Tissue preparation and histology

Animals were deeply anesthetized and transcardially perfused with 0.1 M phosphate buffer (PB), followed by 4% paraformaldehyde (PFA) in 0.1 M PB (pH 7.4). Brains were removed and post-fixed in 4% PFA at 4°C for 48 hours. After cryoprotection, the brains were embedded in OCT compound and sectioned at 40 µm thickness using a Leica CM3050 S cryostat (Leica Microsystems, Buffalo Grove, IL). Sections were permeabilized and blocked for 1 hour at room temperature in PB containing 0.1% Triton X-100 and 10% normal goat serum (NGS). They were then incubated overnight at 4°C with a rabbit anti-GFP primary antibody (Invitrogen; 1:1000 dilution) prepared in PB containing 10% NGS. After three washes in PB, sections were incubated for 1 hour at room temperature with a goat anti-rabbit IgG secondary antibody conjugated to Alexa Fluor 488 (Invitrogen; 1:500 dilution) in PB containing 0.5% NGS. Following another three washes, the sections were mounted on microscope slides (Brain Research Laboratories, Waban, MA) using DAPI Fluoromount-G mounting medium (Southern Biotech) and coverslipped. Coronal brain slices were collected at specific bregma levels to examine the anterior-posterior distribution of the cannula location and viral spread. For ACA analysis, slices were taken at +0.62, +0.26, and −0.22 mm from bregma, based on the Allen Brain Atlas. For visual cortex, slices were collected at −3.40, −3.80, and −4.24 mm. Basal forebrain slices were selected at +0.14, −0.22, and −0.94 mm. To assess viral expression and cannula placement, fluorescent images were acquired using an EVOS FL Imaging System (Thermo Fisher Scientific, Waltham, MA). Imaging was performed with a 10× objective lens for both GFP and DAPI channels. To ensure consistency across imaging sessions, GFP signal was captured with the LED light source set to 90% intensity, while DAPI images were taken at 20–30% intensity. Representative high-resolution images were obtained using an inverted confocal microscope (LSM 780, Zeiss), also with a 10× objective lens. Image analysis was performed using ImageJ software (National Institutes of Health). To analyze optic fiber placement, cannula tracks were mapped onto coronal brain templates based on the Paxinos and Franklin mouse brain atlas. The lowest point of the drawn oval indicates the depth at which the cannula tip was positioned within the tissue.

#### Statistical methods

All statistical analyses were performed using Prism (GraphPad, CA, USA) or MATLAB (MathWorks, MA, USA). For fiber photometry data, mean fluorescence signals for each trial type were compared using two-way repeated-measures ANOVA (RM ANOVA), with current trial type and past trial type as within-subject factors. Post hoc comparisons were conducted using Tukey’s multiple comparisons test. Correlations between mean signals and correct response latency (Z-score) were also assessed. Area under the receiver operating characteristic curve (auROC) values were calculated using the perfcurve function in MATLAB. For optogenetic experiments, two-way RM ANOVA was used with history and light condition (on vs. off) as within-subject factors, followed by Tukey’s post hoc tests. Paired t-tests were applied for direct comparisons between light-on and light-off conditions when appropriate. One-sample t-tests were conducted to test against baseline or chance-level values. For CRISPR experiments, two-way RM ANOVA followed by Tukey’s post hoc test was used to assess effects of genotype and trial history or past vs. past-past trial types. One-way ANOVA was used when comparing genotype groups alone. For combined CRISPR and fiber photometry experiments, two-way RM ANOVA was used to evaluate interactions between genotype and history, as well as between genotype and current trial type. Proportions of Correct, Incorrect, and Omission trials were analyzed using generalized linear mixed models (GLMMs). Custom scripts were written to assess differences in trial patterns while accounting for subject-level variability. Outliers were detected using Grubbs’ test and excluded from analysis when appropriate. Power analyses were performed using G*Power 3.1 for the optogenetic and CRISPR experiments after preliminary data collection to determine the final sample sizes required to achieve adequate statistical power. For imaging experiments, sample sizes were chosen in accordance with commonly accepted standards in the field. All data are presented as means ± SEM. Behavioral and imaging data were collected from a minimum of 2–5 independently conducted experimental series.

## Acknowledgments

We thank Lundbeck for providing the 15q13.3/+ mouse line, Dr. Peter Rudebeck, Dr. Kazuhisa Shibata, Dr. Atsushi Fujimoto, Dr. Ibuki Harada, and members of the Morishita lab for helpful feedback.

## Funding

National Institute on Aging RF1AG085887 (HM)

National Institute of Mental Health R01MH119523 (HM)

National Institute of Neurological Disorders and Stroke R21NS105119 (HM)

Simons Foundation Autism Research Initiative SFARI 977966 (HM)

Eagles Autism Foundation Award (HM)

National Institute of Mental Health K99MH137314 (TN)

Japan Society for the Promotion of Science Overseas Research Fellowships 202260188 (TN)

National Institutes of Mental Health F31MH121010 (KN)

National Institute on Drug Abuse R01DA054526 (SAk)

## Author contributions

Conceptualization: TN, HM

Methodology: TN, KN, TO, Sak

Validation: TN, SAl, ASe, HL, XW, TO

Formal analysis: TN, JR, YG, AK

Investigation: TN, KN, YG, SAl, ASi, ASe, TM, PM, JB, CC, KC, SL, LW, LB, HL

Resources: YL

Data curation: TN, AK

Writing – original draft: TN

Writing – review & editing: HM

Visualization: TN, YG, AK

Supervision: HM

Project administration: TN, HM

Funding acquisition: TN, HM

## Data and materials availability

Data are available in the main text or the supplementary materials. The rest of relevant data and materials are available from the authors upon reasonable request from the corresponding author.

