## Supplementary material for "Nicotinic modulation of frontal circuits breaks the cycle of attentional performance failure": Fig. S1-S13

### Supplementary figures & legends

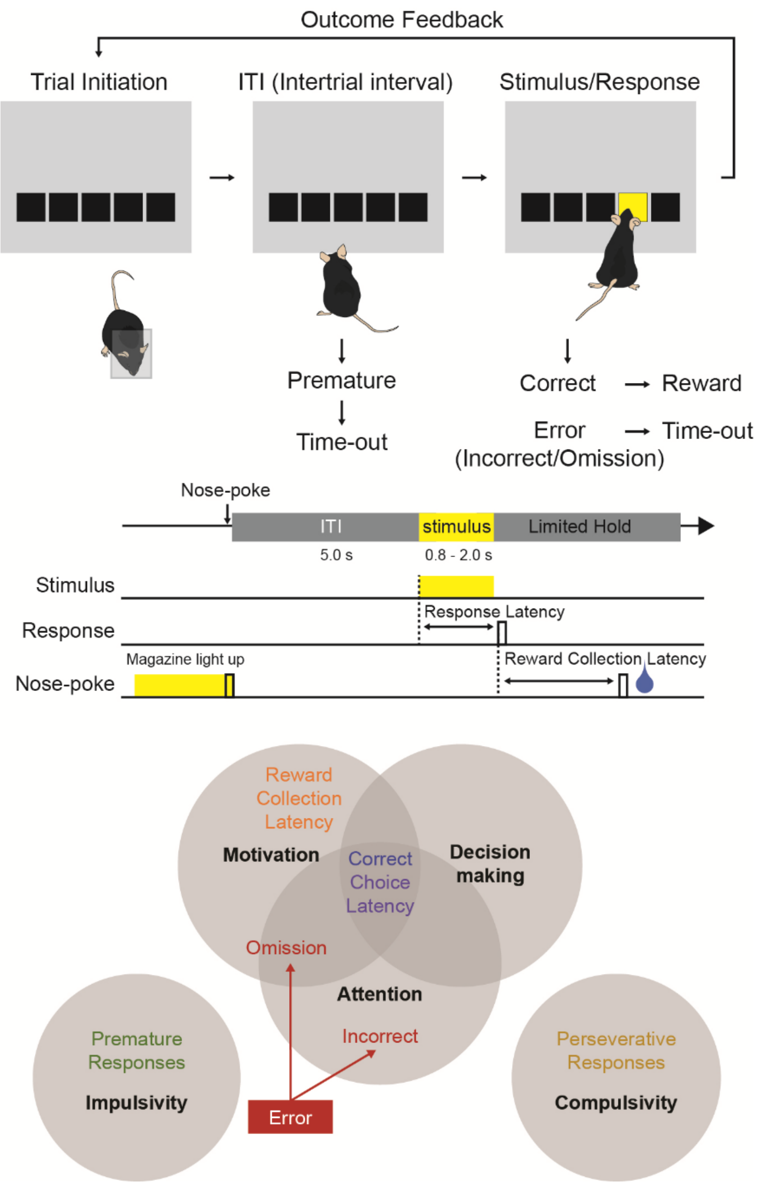


**Fig. S1. Schematic of the 5-choice serial reaction time task (5-CSRTT).** (Top) An overview of 5-CSRTT. 5-CSRTT requires mice to sustain and divide their attentional engagement across five response windows in anticipation of a random presentation of a brief visual stimulus at one of the five locations, and a nose-poke to that location is rewarded. This self-paced, free-moving, attention-demanding task, extensively used in rodent studies, provides a holistic evaluation of a dynamic range of factors that underlie attentional engagement. Mice initiated each trial by nose-poking into the reward magazine. After a 5-s intertrial interval (ITI), a brief visual stimulus (0.8, 1.0, 1.5, or 2.0 s) was randomly presented in one of five apertures. Premature responses during the ITI triggered a 5-s time-out. Correct responses were followed by a chocolate milk reward. Incorrect responses and omissions also resulted in time-outs. Response latency was defined as the time from stimulus onset to aperture response; reward collection latency as the time from correct response to magazine entry. (Bottom) Overview of cognitive functions that contribute to various actions during 5-CSRTT performance. While there is some overlap in measurements used to assess attentional engagement during the 5-CSRTT, each measure has a specific purpose and represents various levels and specificity of attentional performance capacity. Correct response rate is a broad and overall index of attentional performance capacity which reflects what is captured by both accuracy and omission, two readouts which are used to assess attentional engagement level in previous studies using 5-CSRTT. Placing ‘Omissions’ at the intersection of attentional engagement and motivation in the Venn diagram emphasizes that omission probability is sensitive to both. However, another readout of the 5-CSRTT, the latency to collect reward following a correct response, serves as an indicator of motivation to assess if omission change reflects attentional engagement or motivation level. Previous studies have shown that omission rates can change in the absence of corresponding changes in accuracy or reward collection latency, supporting the use of omissions as an index of attentional performance when interpreted in the context of the other task measures *(65)*. Premature responses occurring before stimulus presentation are not included as attentional errors in our analysis because they primarily reflect failures of response inhibition or impulsivity during the inter-trial interval *(19)*.

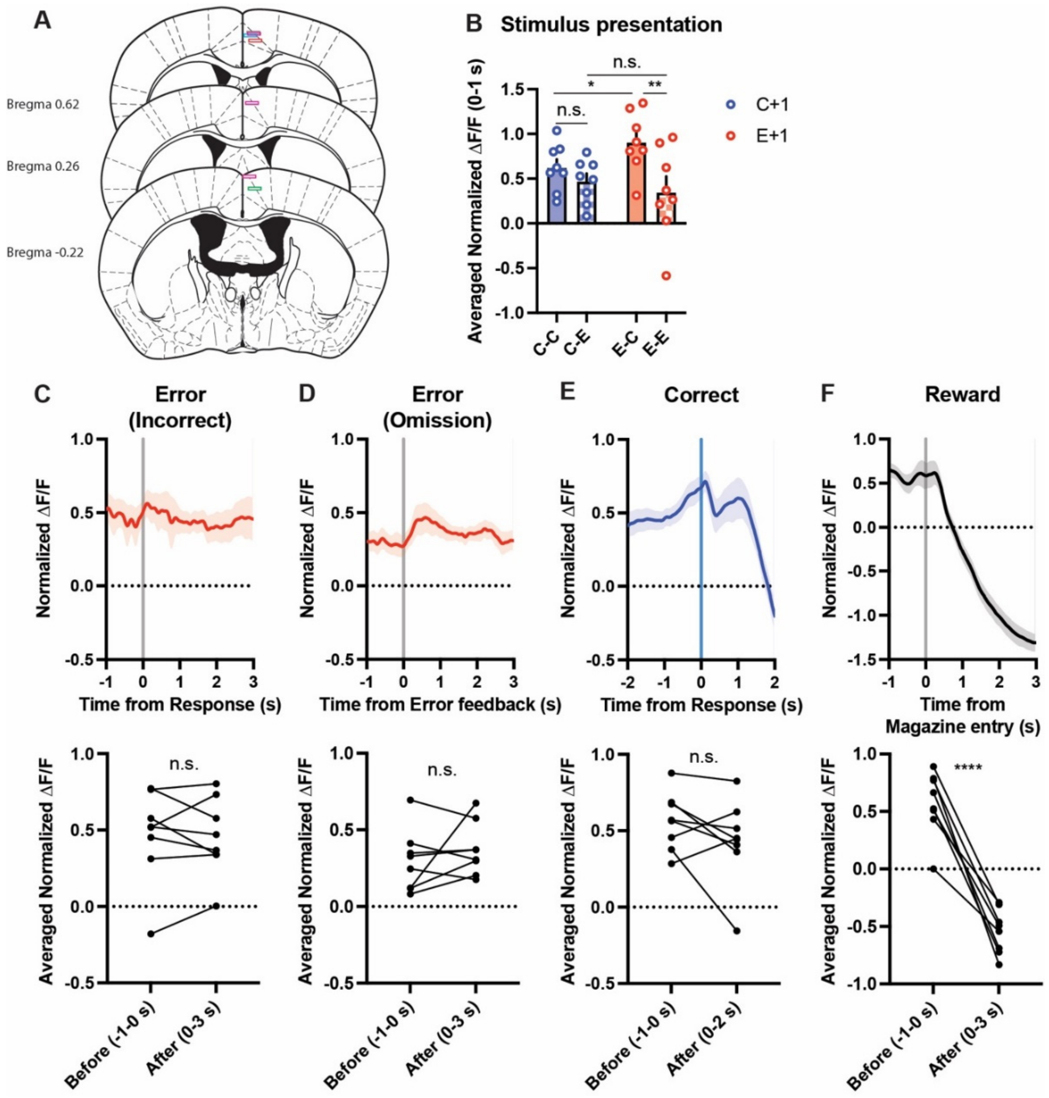


**Fig. S2. ACA$_{VIS}$ activity across trial outcomes in the 5-CSRTT (related to Fig. 1).** (**A**) Histological confirmation of optic fiber placements in the ACA. (**B**) Averaged ACA$_{VIS}$ activity during stimulus presentation (0–1 s). On E+1 trials, activity differed significantly across outcomes (Two-way RM ANOVA, outcome: $F_{1,7} = 61.64$, $P = 0.0001$, n = 8 mice). Post hoc comparisons (Sidak's test) indicated that E-C activity was significantly higher than C-C ($P = 0.040$) and E-E ($P = 0.0012$), whereas C-C vs C-E ($P = 0.28$) or C-E vs E-E ($P = 0.41$) were comparable. (**C–F**) ACA$_{VIS}$ activity aligned to response or feedback: (C) incorrect (Paired t-test, $P = 0.83$), (D) omission (Paired t-test, $P = 0.35$), (E) correct trials (Paired t-test, $P = 0.17$). (F) ACA$_{VIS}$ activity during reward consumption, aligned to magazine entry (Paired t-test, $P < 0.0001$). n.s., not significant; *$P < 0.05$; **$P < 0.01$; ****$P < 0.0001$. Data are mean ± SEM.

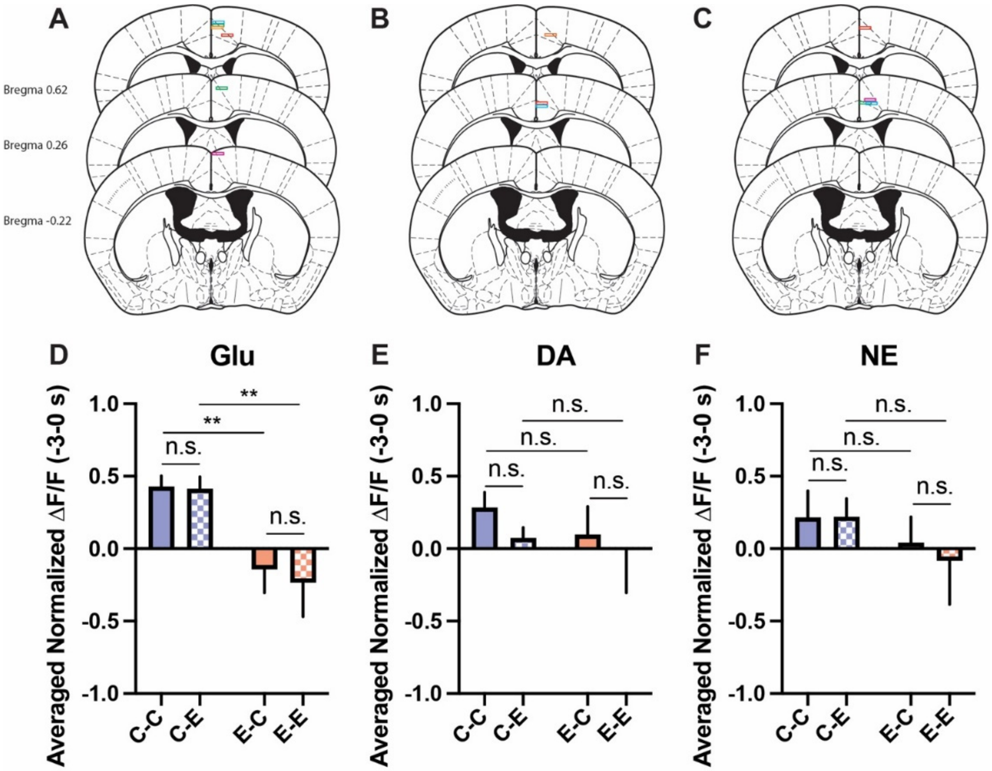


**Fig. S3. Glutamate (Glu), dopamine (DA), and norepinephrine (NE) signals in ACA$_{\mathrm{VIS}}$ across trial outcomes (related to Fig. 1).** (**A–C**) Fiber placements for sensor expression. (A) Glu sensor, (B) DA sensor, and (C) NE sensor. (**D**) Glu activity during the anticipatory period (–3–0 s) differed significantly depending on trial history (Two-way RM ANOVA, effect of history: $F_{1,6} = 12.26$, $P = 0.013$; post hoc Sidak's test: C-C vs. C-E, $P = 0.98$; E-C vs. E-E, $P = 0.62$; C-C vs. E-C, $P = 0.0022$; C-E vs. E-E, $P = 0.0011$; n = 7 mice). (**E**) DA activity did not differ across trial types (Two-way RM ANOVA, effect of history: $F_{1,5} = 0.9842$, $P = 0.37$; post hoc Sidak's test: C-C vs. C-E, $P = 0.11$; E-C vs. E-E, $P = 0.16$; C-C vs. E-C, $P = 0.60$; C-E vs. E-E, $P = 0.63$; n = 6 mice). (**F**) NE activity did not differ between trial types (Two-way RM ANOVA, effect of history: $F_{1,3} = 1.886$, $P = 0.26$; post hoc Sidak's test: C-C vs. C-E, $P = 0.99$; E-C vs. E-E, $P = 0.16$; C-C vs. E-C, $P = 0.43$; C-E vs. E-E, $P = 0.16$; n = 4 mice). n.s., not significant; $**P < 0.01$. Data are mean ± SEM.

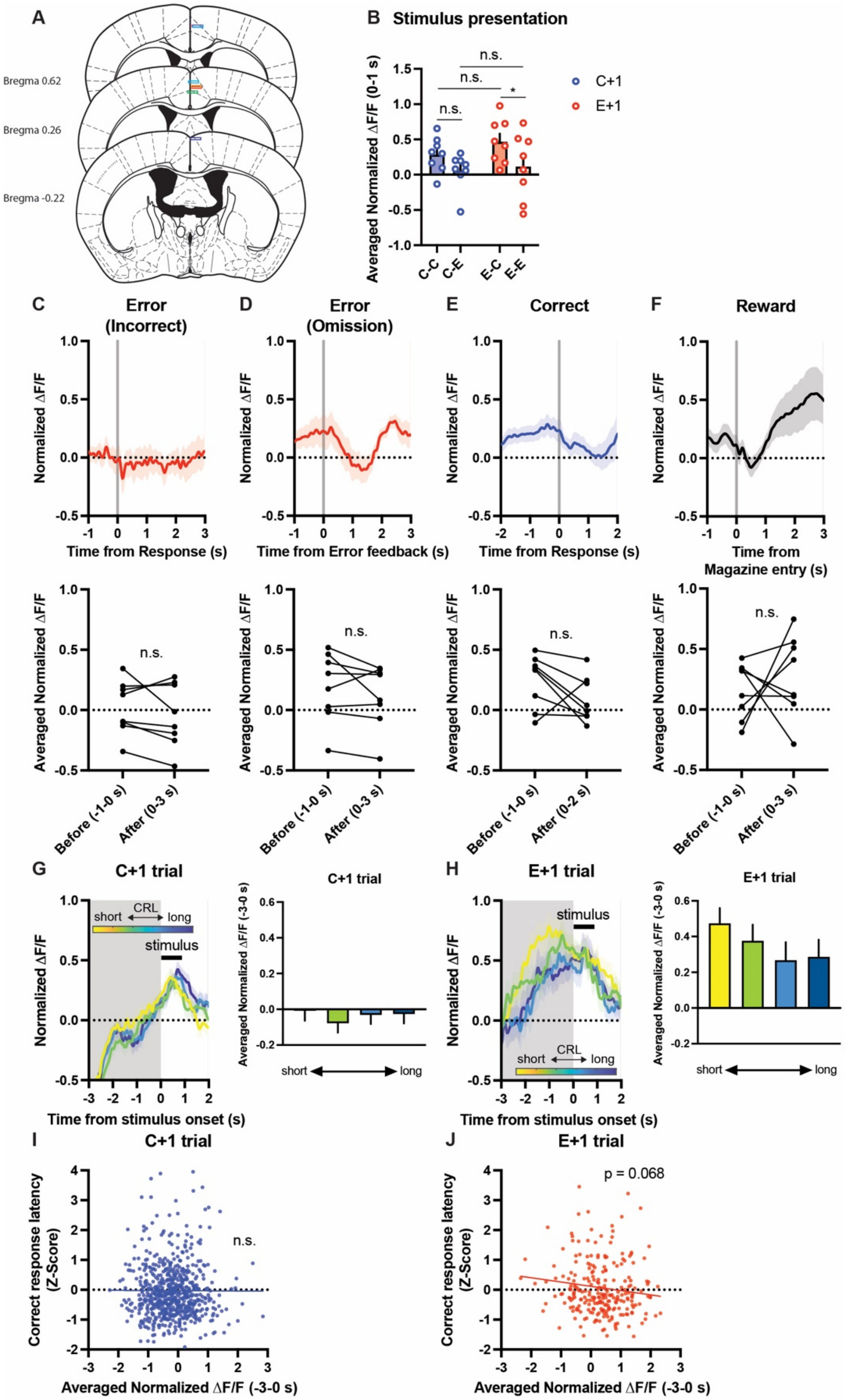


**Fig. S4. Acetylcholine signals in ACA$_{VIS}$ across trial outcomes (related to Fig. 1).** (**A**) Fiber placements for ACh sensor expression. (**B**) Averaged ACh activity during stimulus presentation (0–1 s). Correct responses on E+1 trials showed higher signals than error (Two-way RM ANOVA, effect of current trial outcome: $F_{1,7} = 8.756$, $P = 0.0211$, n = 8 mice). Post hoc comparisons

(Sidak's test) indicated that E-C activity was significantly higher than E-E ($P = 0.016$), whereas C-C vs C-E ($P = 0.093$) or C-E vs E-E ($P = 0.80$) were comparable. (**C–F**) ACh activity aligned to response or feedback: (C) incorrect (Paired t-test, $P = 0.27$), (D) omission (Paired t-test, $P = 0.22$), (E) correct trials (Paired t-test, $P = 0.13$). (F) $ACA_{VIS}$ activity during reward consumption, aligned to magazine entry (Paired t-test, $P = 0.53$). (**G–J**) Relationship between averaged ACh activity and correct response latency. (G, H) Quartile grouping by correct response latency on C+1 and E+1 trials, respectively. (I, J) Trial-by-trial correlation between averaged ACh activity and correct response latency; no significant correlation on C+1 (I, $P = 0.92$), and trend-level correlation on E+1 (J, P = 0.068). n.s., not significant; *$P < 0.05$. Data are mean ± SEM.

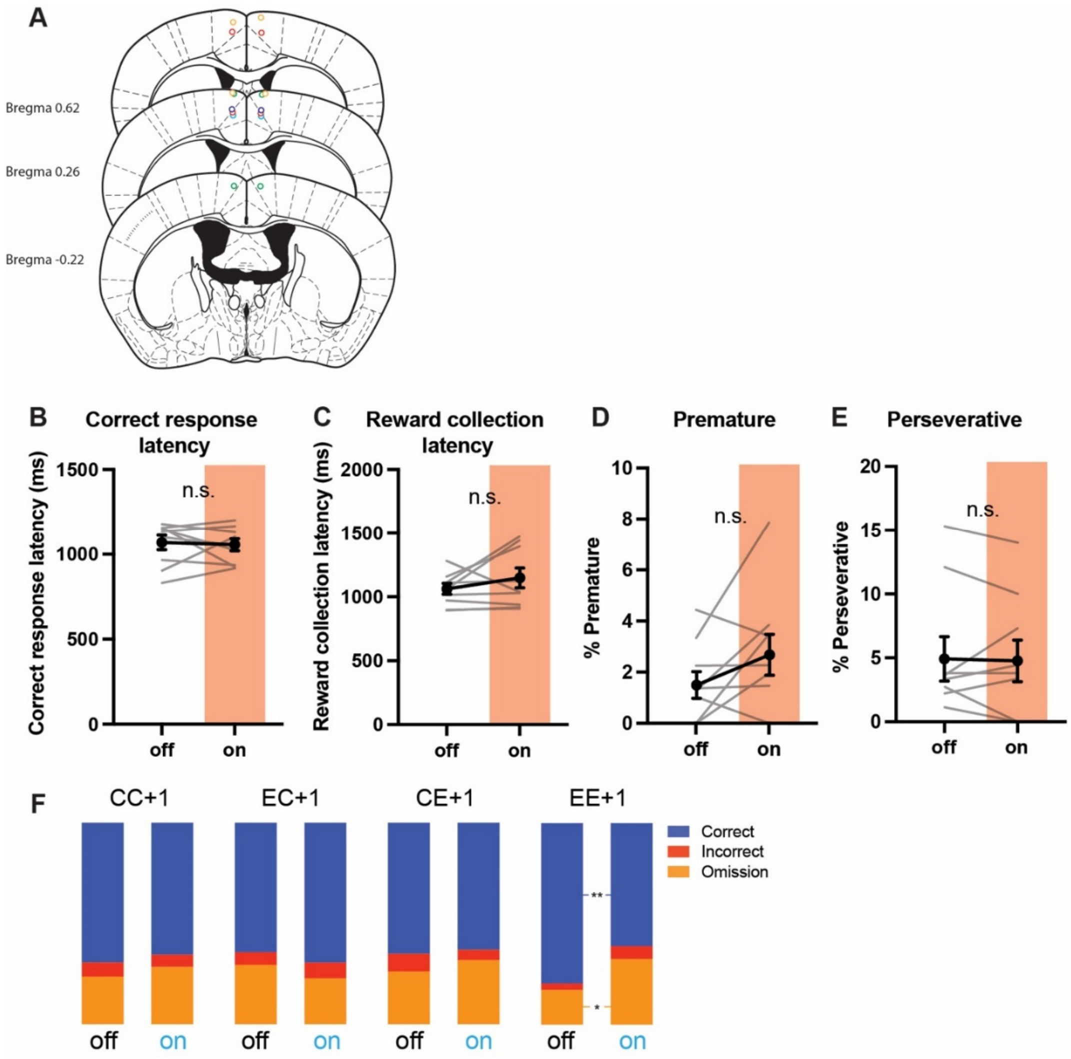


**Fig. S5. Effects of optogenetic inhibition of cholinergic input to ACA on 5-CSRTT performance (related to Fig. 2).** (**A**) Fiber placements for NpHR. (**B–E**) Inhibition had no significant effects on (B) correct response latency (Paired t-test, $P = 0.75$; n = 9 mice), (C) reward collection latency (Paired t-test, $P = 0.24$), (D) premature (Paired t-test, $P = 0.12$), or (E) perseverative responses (Paired t-test, $P = 0.82$). (**F**) Optogenetic inhibition of cholinergic input to ACA significantly increased the percentage of omission on EE+1 trials (Generalized Linear Mixed Model [GLMM], $P = 0.026$). n.s., not significant; *$P < 0.05$; **$P < 0.01$. Data are mean ± SEM.

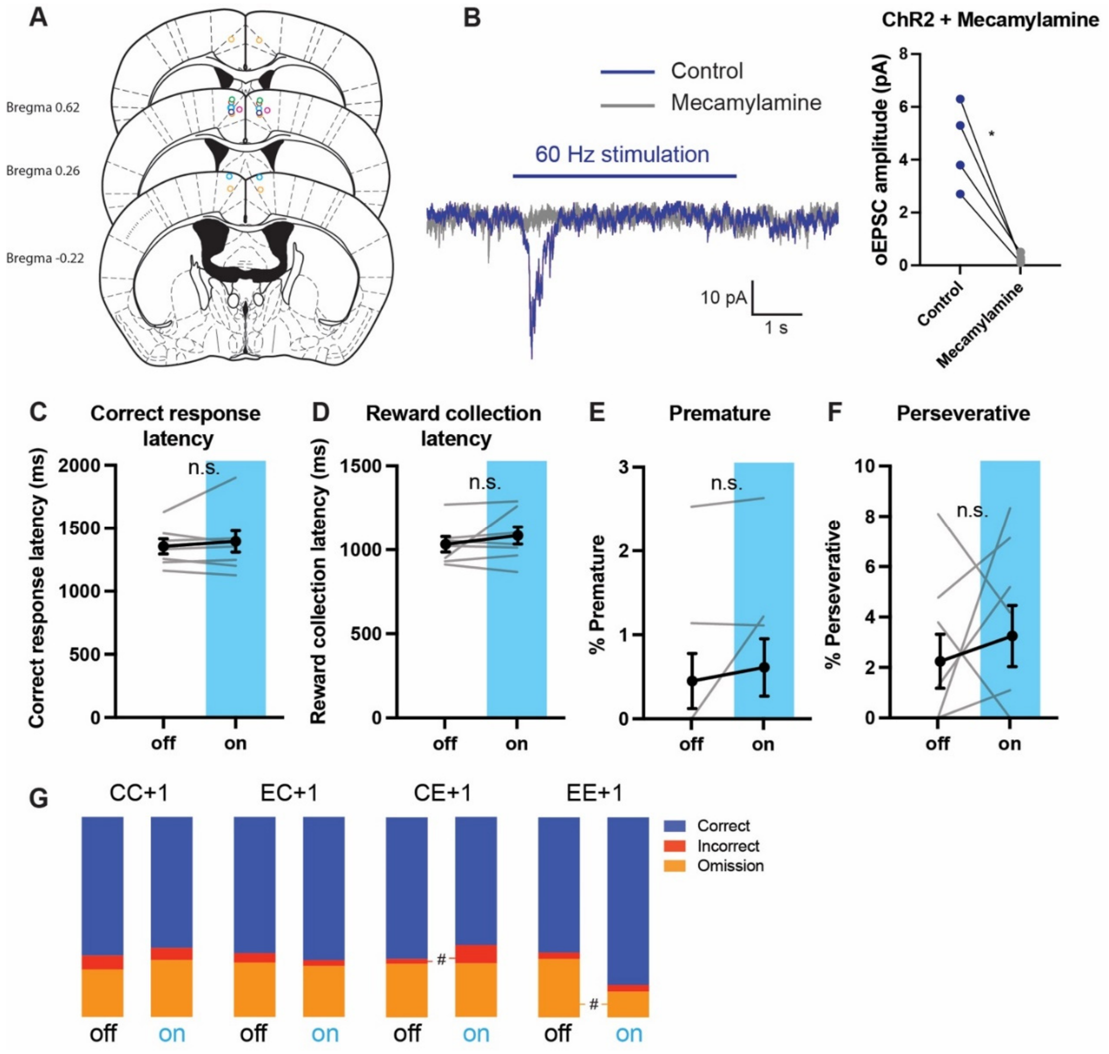


**Fig. S6. Effects of optogenetic activation of cholinergic input to ACA on 5-CSRTT performance (related to Fig. 2).** (**A**) Fiber placements for ChR2. (**B**) Photostimulation of basal forebrain (BF) axon terminals at 60 Hz evokes ACh-mediated inward currents in $ACA_{VIS}$ neurons. Left: representative trace. Right: average optogenetically evoked excitatory postsynaptic current (oEPSC), which was abolished by the ACh receptor antagonist mecamylamine (Paired t test, $P = 0.013$, n = 4 cells from 3 mice). (**C–F**) Activation had no significant effects on (C) correct response latency (Paired t test, $P = 0.61$; n = 8 mice), (D) reward collection latency (Paired t test, $P = 0.35$), (E) premature (Paired t test, $P = 0.32$), or (F) perseverative responses (Paired t test, $P = 0.50$). (**G**) Optogenetic activation of cholinergic input to ACA significantly decreased the percentage of omission on EE+1 trials (Generalized Linear Mixed Model [GLMM], $P = 0.083$). n.s., not significant; #$P < 0.10$; *$P < 0.05$; **$P < 0.01$. Data are mean ± SEM.

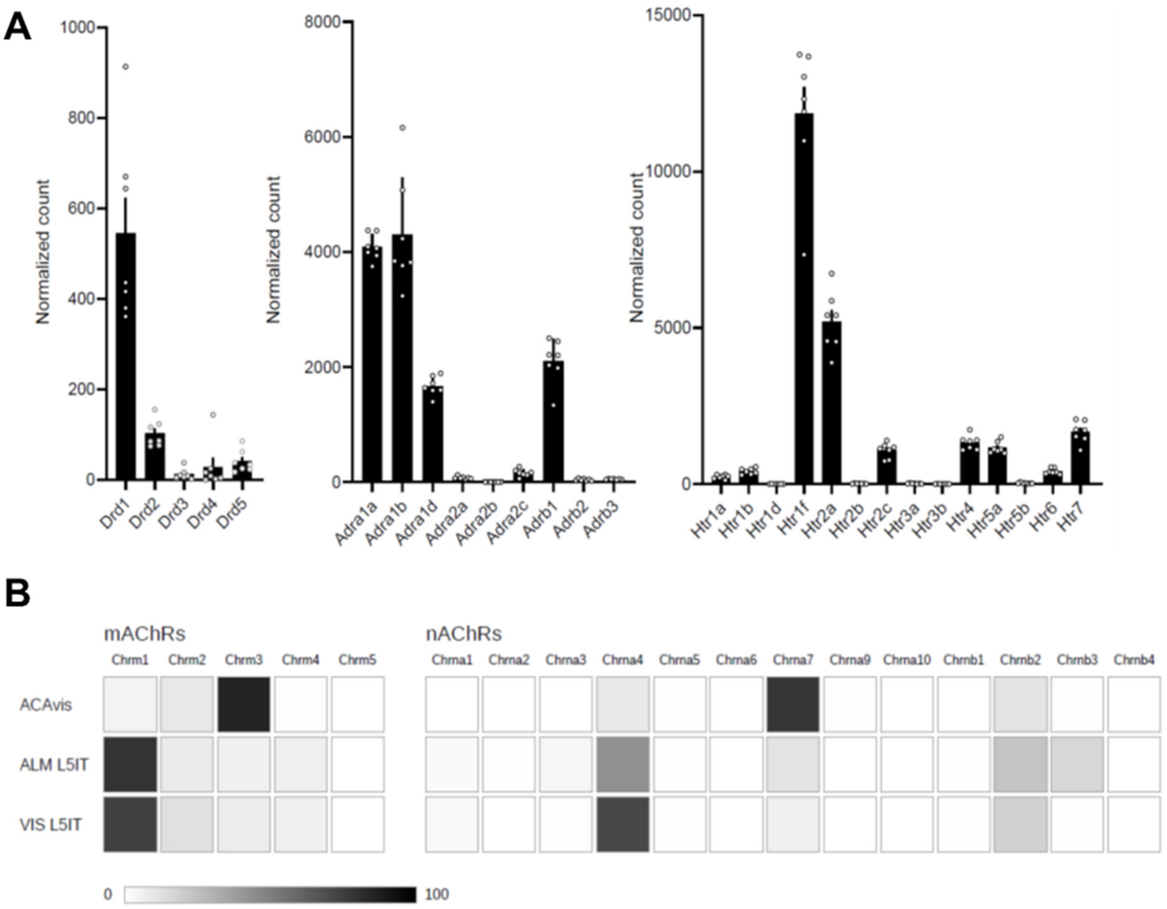


**Fig. S7. Expression profile of neuromodulatory receptors in ACAvis neurons (related to Fig. 3). A.** Expression profile of dopaminergic (Drd), adrenergic (Adra), and serotonergic (Htr) receptor subtypes in ACAvis neurons, based on normalized RNA-seq read counts using DESeq2 size-factor normalization. **B.** Expression profiles of mAChRs and nAChRs in ACAvis (based on Fig.3B), ALM L5IT, and VIS L5IT cell-types (from Allen Institute cell types database: V1 & ALM - SMART-seq, 2018). The relative expression levels of receptor subtypes within each cell type are represented by a grayscale gradient.

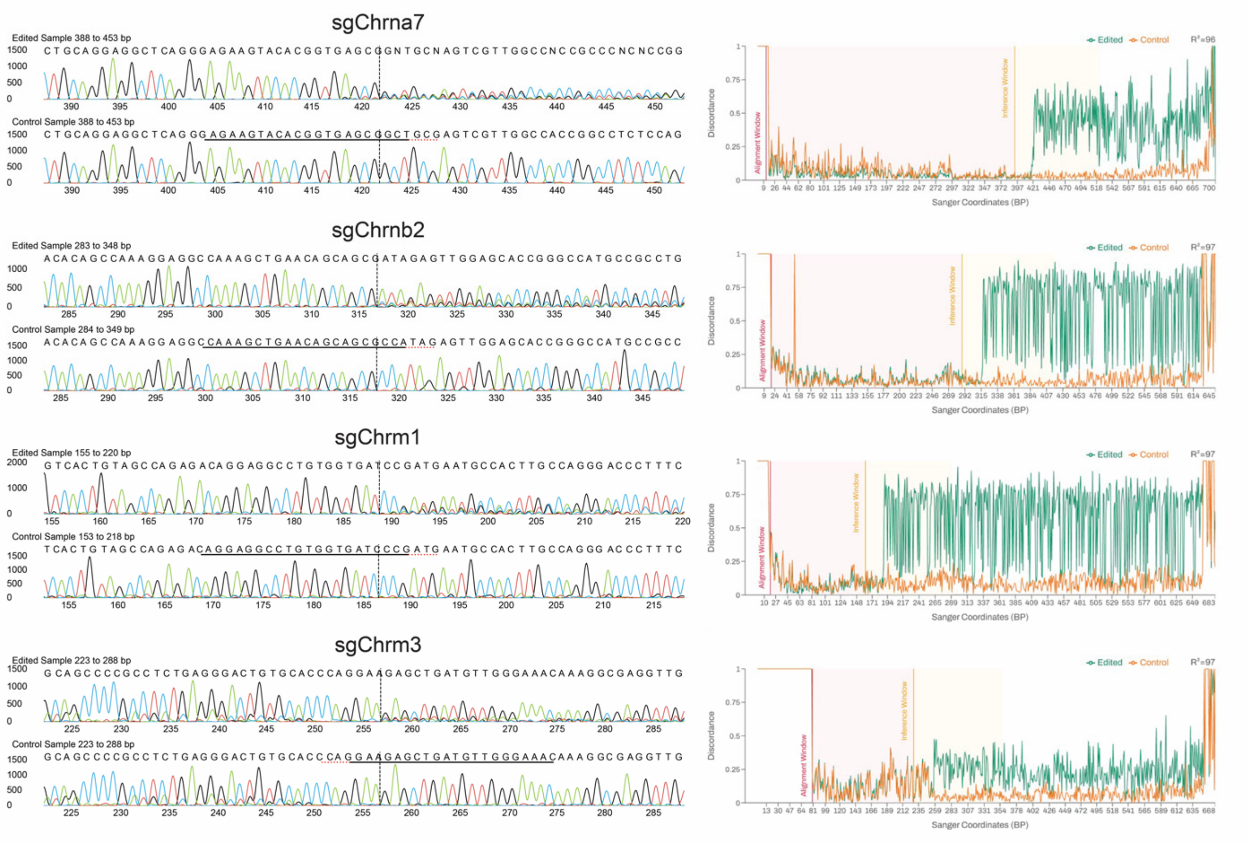


**Fig. S8. Validation of CRISPR-mediated gene editing (related to Fig.3).** *In vitro* validation of CRISPR-mediated gene editing for targeting *Chrna7, Chrnb2, Chrm1, or Chrm3* was performed by sanger sequencing and indel analysis using ICE CRISPR analysis. (Left) The Sanger sequence view showing edited and control sequences in the region around the guide sequence. The horizontal black underlined region represents the guide sequence. The horizontal red underline is the PAM site. The vertical black dotted line represents the actual cut site. Mixed sequencing bases after the cut reflect cutting and error-prone repair by gene editing. (Right) The discordance plots which detail the level of alignment per base between the control and the edited sample in the inference window. It shows the average amount of signal that disagrees with the reference sequence derived from the control trace. For all panels, the green line and orange line are close together before the cut site, with CRISPR edits resulting in a jump in the discordance near the cutsite and continuing to remain far apart after the cut site.

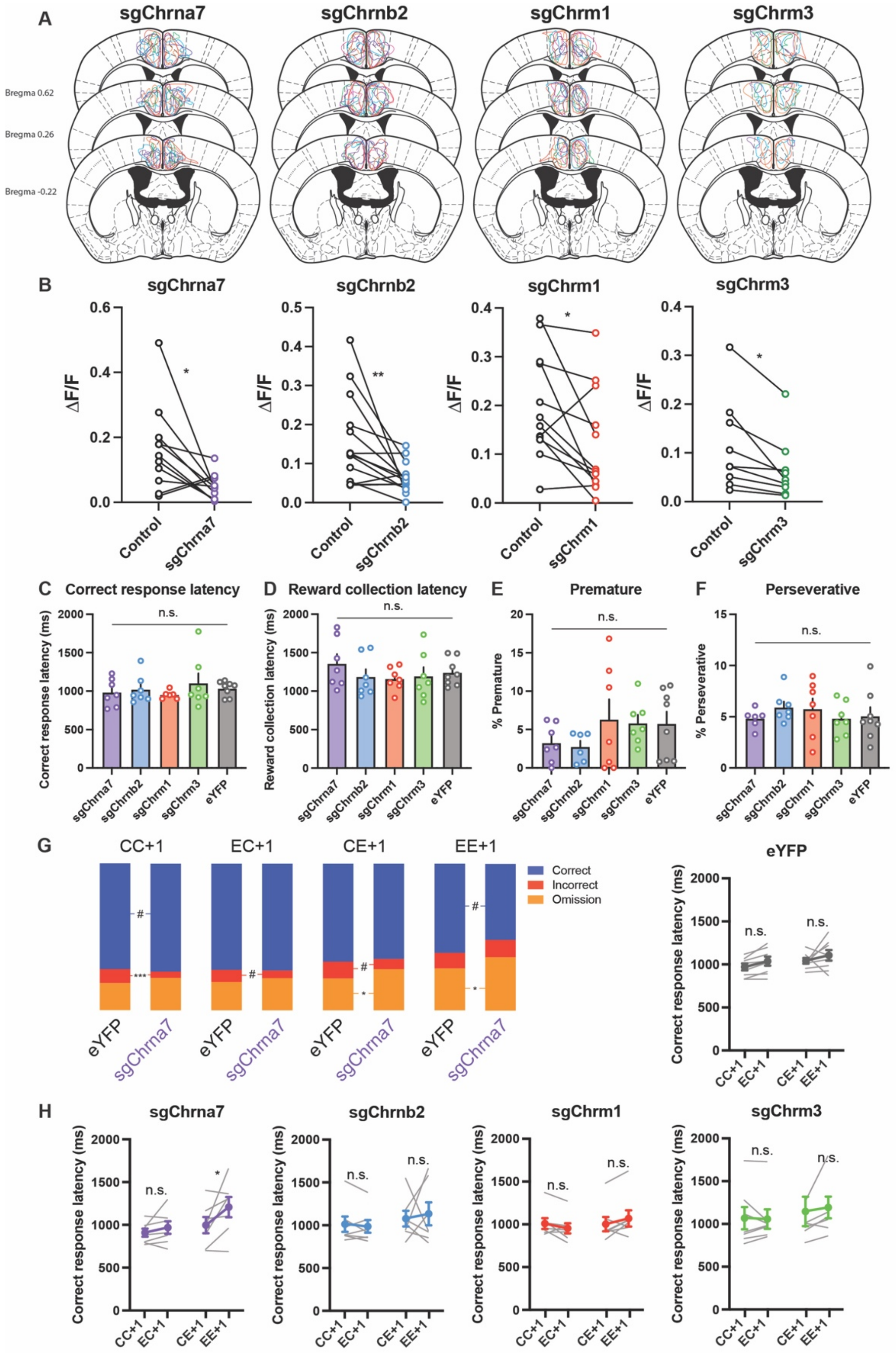


**Fig. S9. Effects of AChR subtype knockdown in $ACA_{VIS}$ on 5-CSRTT performance (related to Fig. 3).** (**A**) AAV spread summary for each knockdown: sgChrna7, sgChrnb2, sgChrm1, sgChrm3. Viral spread of individual animals are shown using different colors for each case. (**B**) Functional validation of CRISPR AAV: Expression of sgRNAs significantly attenuated cholinomimetic carbachol-evoked responses (Paired t-test: sgChrna7, $P = 0.021$, n = 11 slices from

4 mice; sgChrnb2, $P = 0.0098$, n = 14 slices from 5 mice; sgChrm1, $P = 0.021$, n = 12 slices from 4 mice; sgChrm3, $P = 0.011$, n = 9 slices from 4 mice). (**C-F**) Knockdown did not affect (C) correct response latency (One-way ANOVA, $P = 0.69$; n = 7 mice for sgChrna7, sgChrnb2, sgChrm1, and sgChrm3; n = 8 for eYFP), (D) reward collection latency (One-way ANOVA, $P = 0.61$), (E) premature (One-way ANOVA, $P = 0.41$), or (F) perseverative responses (One-way ANOVA, $P = 0.76$). (**G**) *Chrna7* knockdown significantly increased the percentage of omission errors on EE+1 trials (Generalized Linear Mixed Model [GLMM], $P = 0.045$). **(H)** *Chrna7* knockdown increased correct response latency specifically on EE+1 trials (Two-way RM ANOVA, past × past-past trial: $F_{1,12} = 6.029$, $P = 0.030$; post hoc Sidak's test: CC+1 vs. EC+1, $P = 0.71$; EE+1 vs. CE+1, $P = 0.038$). No such effects were observed in other knockdowns or in the control group (post hoc Sidak's test: sgChrnb2, CC+1 vs. EC+1, $P = 0.97$, EE+1 vs. CE+1, $P = 0.88$; sgChrm1, CC+1 vs. EC+1, $P = 0.37$, EE+1 vs. CE+1, $P = 0.24$; sgChrm3, CC+1 vs. EC+1, $P = 0.99$, EE+1 vs. CE+1, $P = 0.87$; eYFP, CC+1 vs. EC+1, $P = 0.35$, EE+1 vs. CE+1, $P = 0.38$). n.s., not significant; #$P < 0.10$; *$P < 0.05$; **$P < 0.01$; ***$P < 0.001$. Data are mean ± SEM.

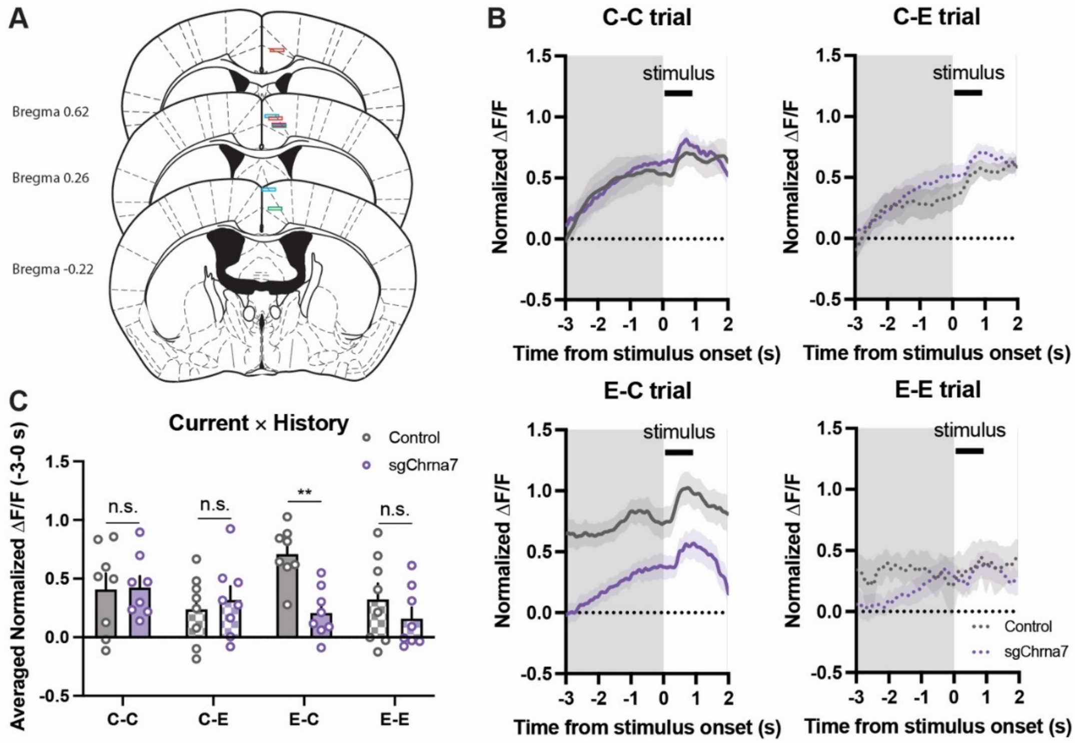


**Fig. S10. ACA$_{VIS}$ neuron activity following α7-nAChR knockdown during the 5-CSRTT (related to Fig. 4).** (**A**) Schematic of fiber placement for jGCaMP8s imaging. (**B**) Neural activity dynamics in ACA$_{VIS}$ neurons across different trial history conditions (C–C, C–E, E–C, E–E) in control and sgChrna7 mice. (**C**) Averaged normalized ΔF/F during the anticipatory attentional engagement period (–3 to 0 s before stimulus onset), grouped by past and current trial outcomes (C–C, C–E, E–C, E–E), for control and sgChrna7 mice. sgChrna7 mice showed reduced ACA$_{VIS}$ activity in E–C trials (Two-way RM ANOVA, genotype × trial-type interaction: $F_{3,42}$ = 4.727, $P$ = 0.0063; post hoc Sidak's test: C-C, $P$ > 0.99; C-E, $P$ = 0.97; E-C, $P$ = 0.0047; E-E, $P$ = 0.72; n = 8 mice per group). n.s., not significant; **$P$ < 0.01. Data are mean ± SEM.

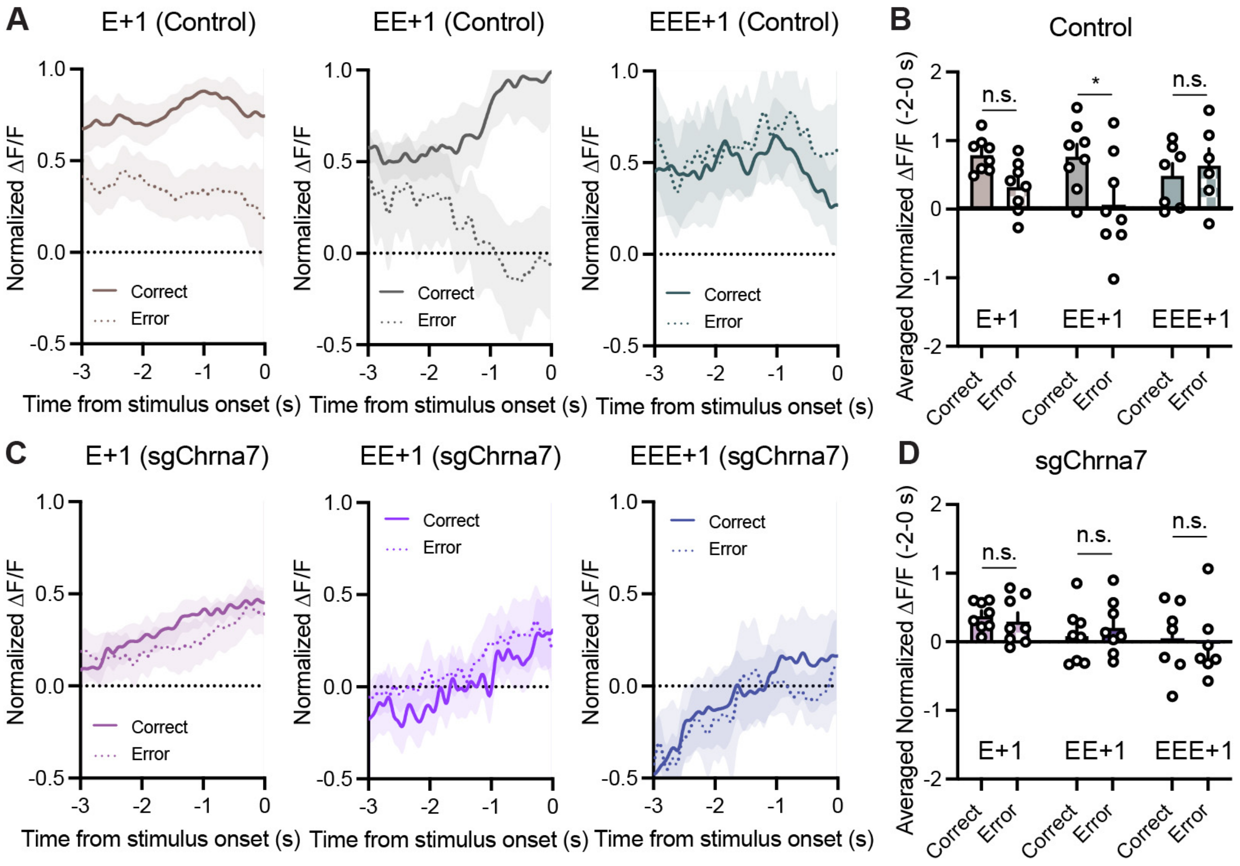


**Fig. S11. Impact of the number of consecutive prior errors on the ability of ACAvis activity to predict behavioral performance during the 5-CSRTT (related to Fig. 4).** (**A**) Neural activity in ACAvis during the anticipatory attentional engagement period in control mice following a single error (E+1, left), two consecutive errors (EE+1, middle), and three consecutive errors (EEE+1, right). (**B**) In control mice, ACAvis activity on EE+1 trials was significantly higher in correct than in error trials (Two-way RM ANOVA, trial type factor: $F_{1,18} = 6.631$, $P = 0.019$; post hoc Sidak's test: E+1, $P = 0.14$; EE+1, $P = 0.016$; EEE+1, $P = 0.93$). (**C**) Neural activity in ACAvis during the anticipatory attentional engagement period in sgChrna7 mice following a single error (E+1, left), two consecutive errors (EE+1, middle), and three consecutive errors (EEE+1, right). (**D**) In sgChrna7 mice, ACAvis activity did not differ between correct and error trials for any trial type (Two-way RM ANOVA, trial type factor: $F_{1,40} = 0.05286$, $P = 0.82$; post hoc Sidak's test: E+1, $P = 0.95$; EE+1, $P = 0.95$; EEE+1, $P = 0.98$). n.s., not significant; *$P < 0.05$; **$P < 0.01$. Data are presented as mean ± SEM.

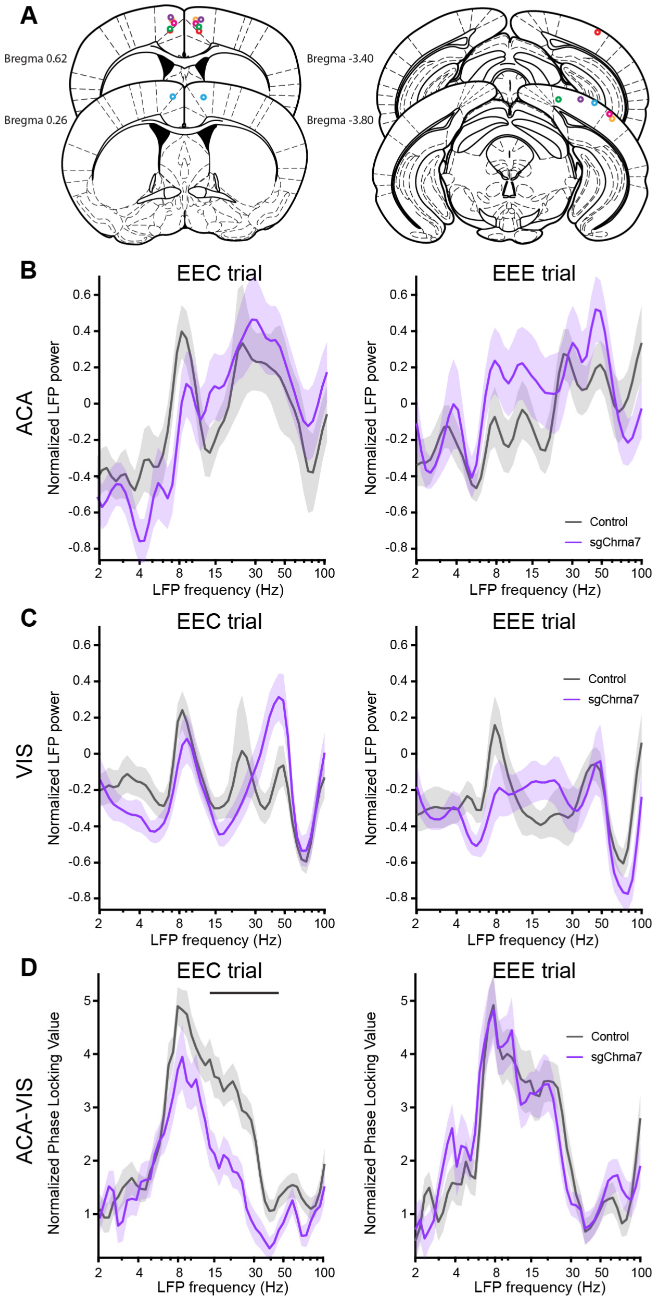


**Fig. S12. LFP data following repeated errors during 5-CSRTT in control and sgChrna7 mice (related to Fig. 4).** (**A**) Schematic of electrode placement for LFP recording. (**B, C**) Normalized LFP power across frequency range in ACA (B) and VIS (C) in control and sgChrna7 mice during anticipatory attentional engagement period (−3 to 0 s before stimulus onset) on EEC (left) or EEE trials (right). (**D**) Normalized phase-locking value (PLV) between ACA and VIS across the frequency range in control and sgChrna7 mice during the anticipatory attentional engagement period on EEC (left) or EEC trials (right**)**. The black line indicates frequency clusters in which the metric differs across genotypes ($P < 0.05$, cluster-based permutation test). Low-gamma PLV is significantly lower in sgChrna7 mice compared to control mice on EEC trials but not on EEE trials. n = 3 mice per group.

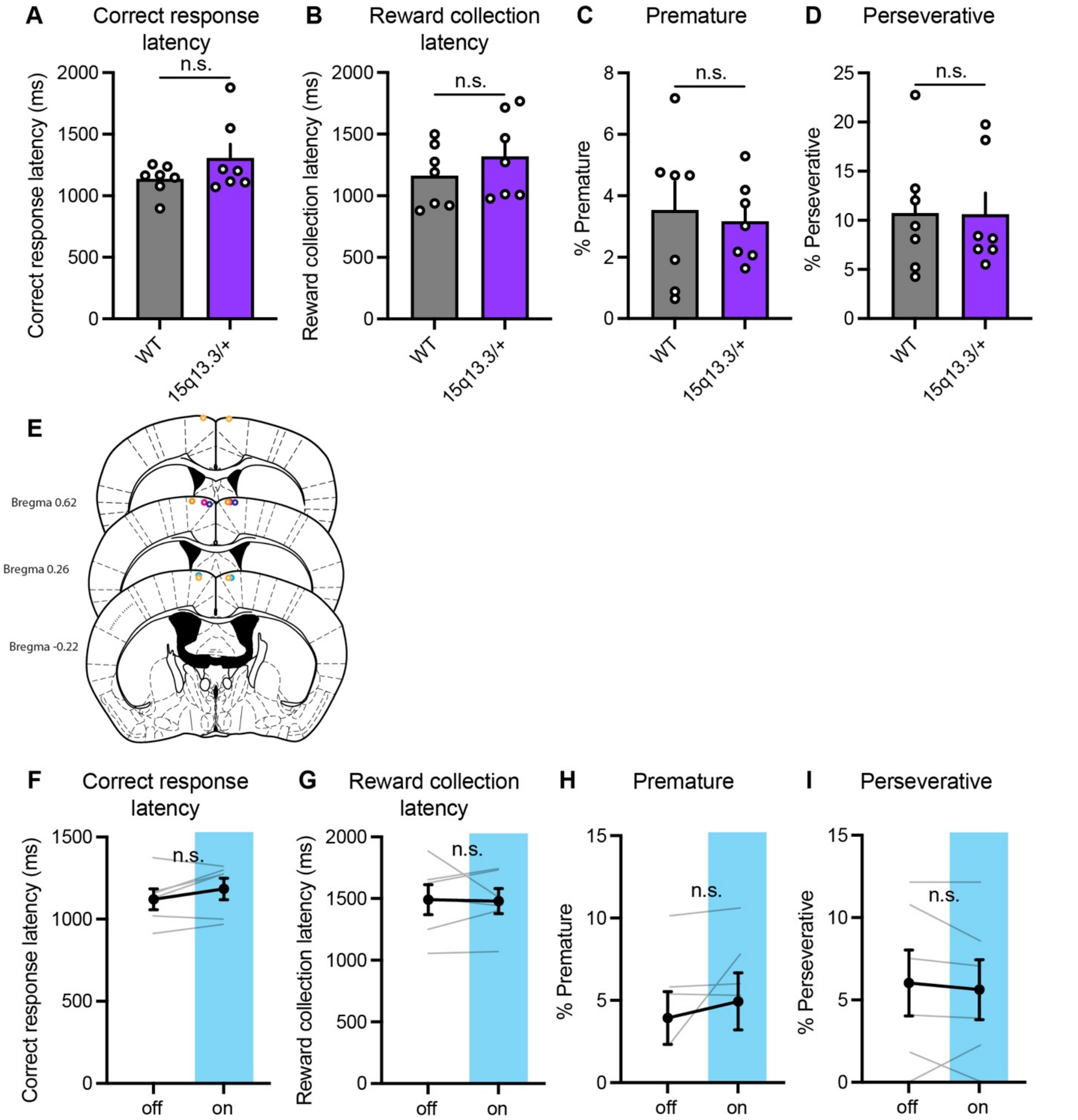


**Fig. S13. Comparable performance between WT and 15q13.3/+ mice in non-attentional measures of the 5-CSRTT, unaffected by optogenetic activation of ACA neurons (related to Fig. 5).** (**A–D**) No significant differences were observed between WT and 15q13.3/+ mice in (A) correct response latency (Unpaired t-test, $P = 0.20$, n = 7 mice per group), (B) reward collection latency (Unpaired t test, $P = 0.35$), (C) premature responses (Unpaired t test, $P = 0.73$), or (D) perseverative responses (Unpaired t test, $P = 0.97$). (**E**) Fiber placements for ChR2. (**F–I**) Optogenetic activation had no significant effects on (C) correct response latency (Paired t test, $P = 0.13$; n = 6 mice), (D) reward collection latency (Paired t test, $P = 0.89$), (E) premature (Paired t test, $P = 0.31$), or (F) perseverative responses (Paired t test, $P = 0.56$). n.s., not significant. Data are mean ± SEM.
